# Staged maturation converts a probasal body template complex into a nucleation-competent basal plate

**DOI:** 10.64898/2026.09.23.753794

**Authors:** Athina Paterou, Samuel Dean

## Abstract

The central pair (CP) of microtubules is essential for flagellar motility, yet the molecular events underpinning its nucleation remain poorly defined. Here, we report a comprehensive screen of flagellar transition zone proteins that identifies new central pair assembly proteins (CPAPs). Using expansion microscopy, we reveal a concentric architecture at the basal plate, with themelin and basalin forming nested punctae with radial symmetry surrounding a central core containing TZP103.8, TZP48 and γ-tubulin. A subset of CPAPs pre-assemble at the probasal body, a precursor structure that has not yet built a flagellum, with themelin functioning as the architectural template and master recruiter, and TZP103.8 and γ-tubulin IFT-dependant addition being the final step of basal plate activation. Strikingly, in the absence of TZP103.8, γ-tubulin still reaches the basal plate but is insufficient for central pair nucleation. We propose the staged assembly of CPAPs progressively constructs a structural template that geometrically constrains γ-tubulin into the positions and orientations required for CP nucleation.

## Introduction

Motile cilia and flagella are ancient organelles that were present in the last eukaryotic common ancestor and remain essential across the tree of life (1). In single-celled organisms, flagella provide cellular propulsion (2), while in metazoa they also drive the movement of fluids across ciliated epithelia (3). In humans, defects in flagellar motility cause primary ciliary dyskinesia (PCD), a condition estimated to affect 1 in 10,000–40,000 people, resulting in chronic respiratory infections, laterality defects and reduced fertility (4). In kinetoplastid parasites, including *Trypanosoma brucei*, the causative agent of African sleeping sickness, flagellar motility is essential for infectivity and transmission via the tsetse fly vector (5–7).

The core of the motile flagellum is the axoneme, a highly conserved microtubule-based scaffold characterised by nine outer microtubule doublets surrounding a central pair (CP) of singlet microtubules - the so-called 9+2 arrangement (8, 9). Flagellar motility is driven by dynein motor complexes attached along the outer doublets (9), whose activity is coordinated in part by the CP via radial spoke connections (10, 11). Loss of the CP, or of CP-associated projections, leads to flagellar paralysis and is associated with PCD in humans (12, 13). Despite the central importance of the CP to flagellar motility, the molecular mechanisms governing its nucleation and assembly remain poorly understood.

The CP microtubules are nucleated at the distal end of the transition zone (TZ), a structurally and functionally distinct region at the base of the flagellum that acts as a diffusion barrier and gate controlling ciliary composition (14, 15). At the TZ–axoneme boundary, an electron-dense structure called the basal plate serves as the site of CP nucleation (16, 17). Previously, we identified two components of the basal plate in *T. brucei* that are essential for CP assembly: basalin and TZP103.8 (16, 18). Ablation of basalin prevented assembly of the basal plate itself, indicating that it functions as a core ‘hub’ protein onto which other basal plate components are recruited. In contrast, loss of TZP103.8 abolished CP nucleation but left the basal plate structurally intact, placing TZP103.8 functionally downstream of basalin in the basal plate and CP assembly pathway. γ-tubulin, which templates the nucleation of new microtubule protofilaments throughout the cell, is also required for CP formation (19). However, fluorescence microscopy has not detected γ-tubulin at the basal plate; instead, the dominant signal localises to the proximal end of the basal body and probasal body (16, 20). Moreover, other components of the γ-tubulin small complex (γTuSC) have not been reported at the basal plate (21), raising the question of whether γ-tubulin acts directly at this site or via an unconventional mechanism.

Despite these advances, it remains largely unknown how the central pair is nucleated such that exactly two microtubules form at defined positions and orientations, and at the correct developmental stage.

Here, we report a comprehensive RNAi screen of TZ components in *T. brucei* that identifies two additional proteins required for CP assembly. Using ultrastructure expansion microscopy (U-ExM) and iterative U-ExM (iU-ExM), we resolve the molecular architecture of the basal plate and discover that a subset of CP assembly factors form an immature template at the probasal body - a precursor structure that has not yet nucleated a flagellum - establishing that preparation for CP nucleation begins far earlier in basal body ontogenesis than previously appreciated. Finally, we reveal a previously unsuspected role for IFT in ‘activating’ the basal plate for CP nucleation.

## Results

### A targeted RNAi screen reveals two new central pair assembly proteins

To identify additional proteins required for CP assembly, we developed a targeted RNAi screen exploiting a *T. brucei* cell line expressing the CP marker PF16 fused to mScarlet-I as a fluorescent reporter for CP integrity. We serially transfected this reporter line with 44 individual RNAi constructs, each targeting a different transition zone protein (TZP) identified in our earlier proteomics study (18). Following RNAi induction, cells were scored by fluorescence microscopy for loss or disruption of axonemal PF16 signal, indicative of a CP defect (Supplemental Table 1, Fig. 1a).

**Figure 1.**
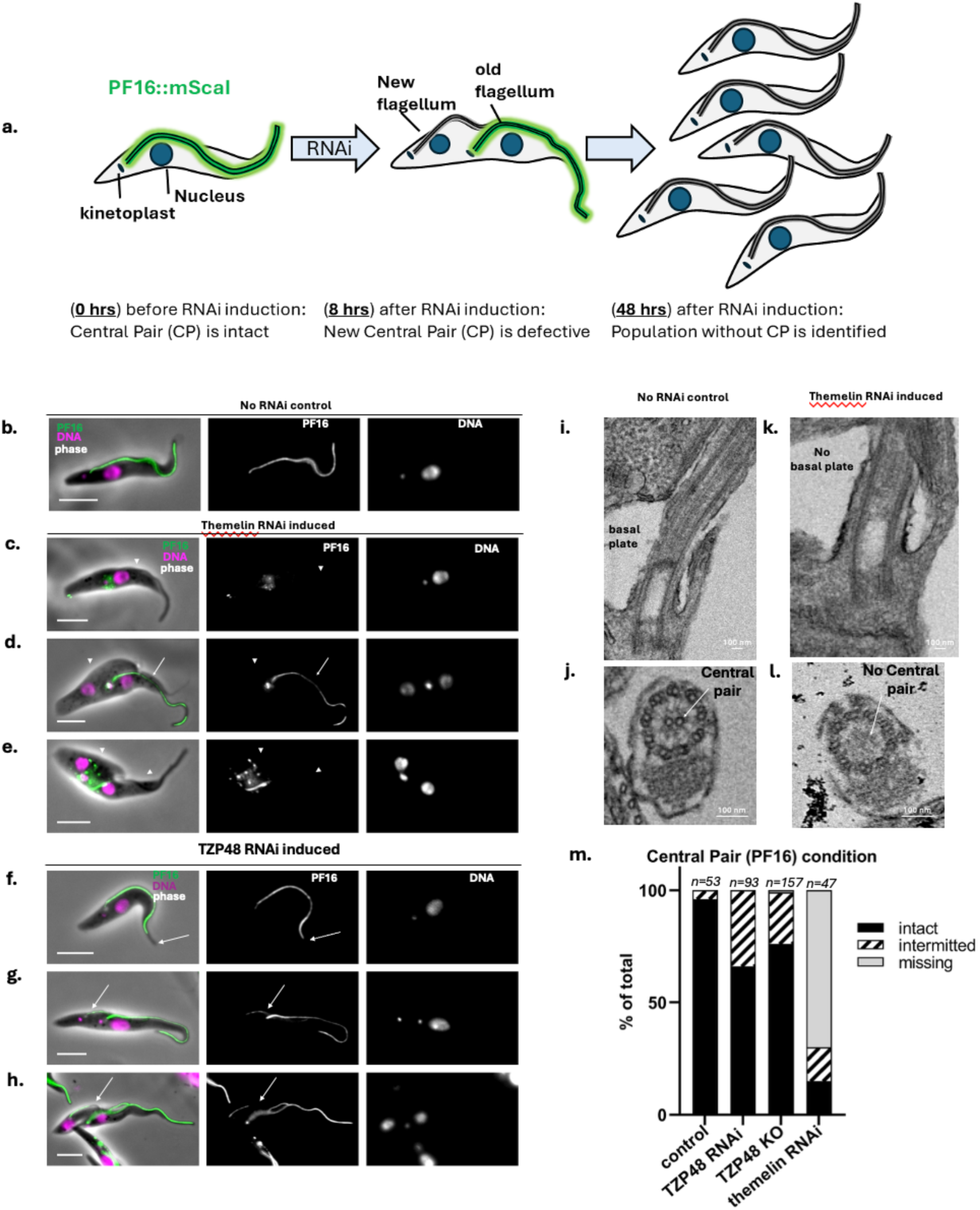
A targeted RNAi screen reveals two new central pair assembly proteins (CPAPs). a) A PF16::mScarlet-I central pair reporter cell line was serially transfected with 48 RNAi constructs, each targeting a different TZP. Cells were induced for RNAi and monitored for missing or intermittent PF16 axonemal fluorescence, indicating a central pair defect. b-h) Representative fluorescence micrographs of control, themelin-ablated cells, or TZP48- ablated cells with absent (arrowheads) or incomplete (arrows) PF16 signal. i-l) TEM longitudinal sections (i, k) and cross sections (j, l) show absence of the central pair and basal plate in themelin ablated cells. m) Categorising PF16 fluorescence in axonemes shows that the themelin RNAi phenotype is more penetrant and severe (48 hrs, 59% missing PF16) than either the TZP48 RNAi (48 hrs, 33% incomplete PF16) or genetic knock out (20% incomplete PF16) phenotypes. Scale bar = 5 μm.

This screen identified two TZPs not previously implicated in CP assembly: TZP46 (Tb927.1.2770) and TZP48 (Tb927.1.4470). TZP46 (46 kDa) has two predicted N-terminal ubiquitin-like domains and a C terminal coiled coil, whereas TZP48 (48 kDa) has no conserved domains detected with high intrinsic disorder (Supplementary Fig. 1). Ablation of TZP46 produced a severe and highly penetrant phenotype, with 59% of axonemes lacking PF16 fluorescence entirely (Fig. 1b–d, j). By comparison, TZP48 ablation yielded a milder phenotype characterised predominantly by weak and intermittent PF16 signal along the axoneme (33% of axonemes affected), rather than complete PF16 loss. To test whether this milder phenotype reflected incomplete knockdown, we generated a TZP48 genetic knockout. This reproduced the intermittent CP phenotype, but at a reduced frequency (20% of axonemes; Fig. 1f–g, j), suggesting that compensatory mechanisms may partially buffer against chronic loss of TZP48. The greater severity of the TZP46 phenotype, and the complete absence rather than mere reduction of PF16, suggested that TZP46 operates upstream of TZP48 in the CP assembly pathway and we therefore prioritised it for further functional analysis.

To determine the ultrastructural basis of the TZP46 ablation phenotype, we examined induced cells by transmission electron microscopy (TEM). Longitudinal sections through the transition zone revealed the absence of the basal plate, while cross-sections through the axoneme confirmed the loss of the CP microtubules (Fig. 1h, i). The loss of the basal plate in TZP46-ablated cells distinguishes the TZP46 phenotype from that of TZP103.8, whose ablation abolishes the CP but leaves the basal plate intact, and instead resembles the basalin ablation phenotype (16, 18). We therefore named this protein themelin (from the Greek θεμέλιο, meaning foundation) and, together with basalin, TZP103.8 and TZP48, designated these four proteins collectively as central pair assembly proteins (CPAPs).

### CPAPs form nested rings surrounding a central core at the basal plate

By standard fluorescence microscopy, each CPAP appears as a diffraction-limited spot at the flagellar base, without sufficient resolution to distinguish its position within the transition zone or basal plate. To resolve their spatial organisation in the TZ, we therefore turned to U-ExM. We combinatorially co-tagged each protein with tandem epitope tags (HA, Myc or FLAG) and stained expanded cells with anti-tag antibodies, alongside anti-alpha tubulin to provide structural context. Cross-sectional and longitudinal views through the basal plate revealed a striking concentric architecture, with themelin occupying the outermost position as a ring that encircled a smaller basalin ring (Fig. 2a, b). In contrast, TZP103.8 and TZP48 each localised to the central core of the basal plate, interior to basalin (Fig. 2c–f).

**Figure 2.**
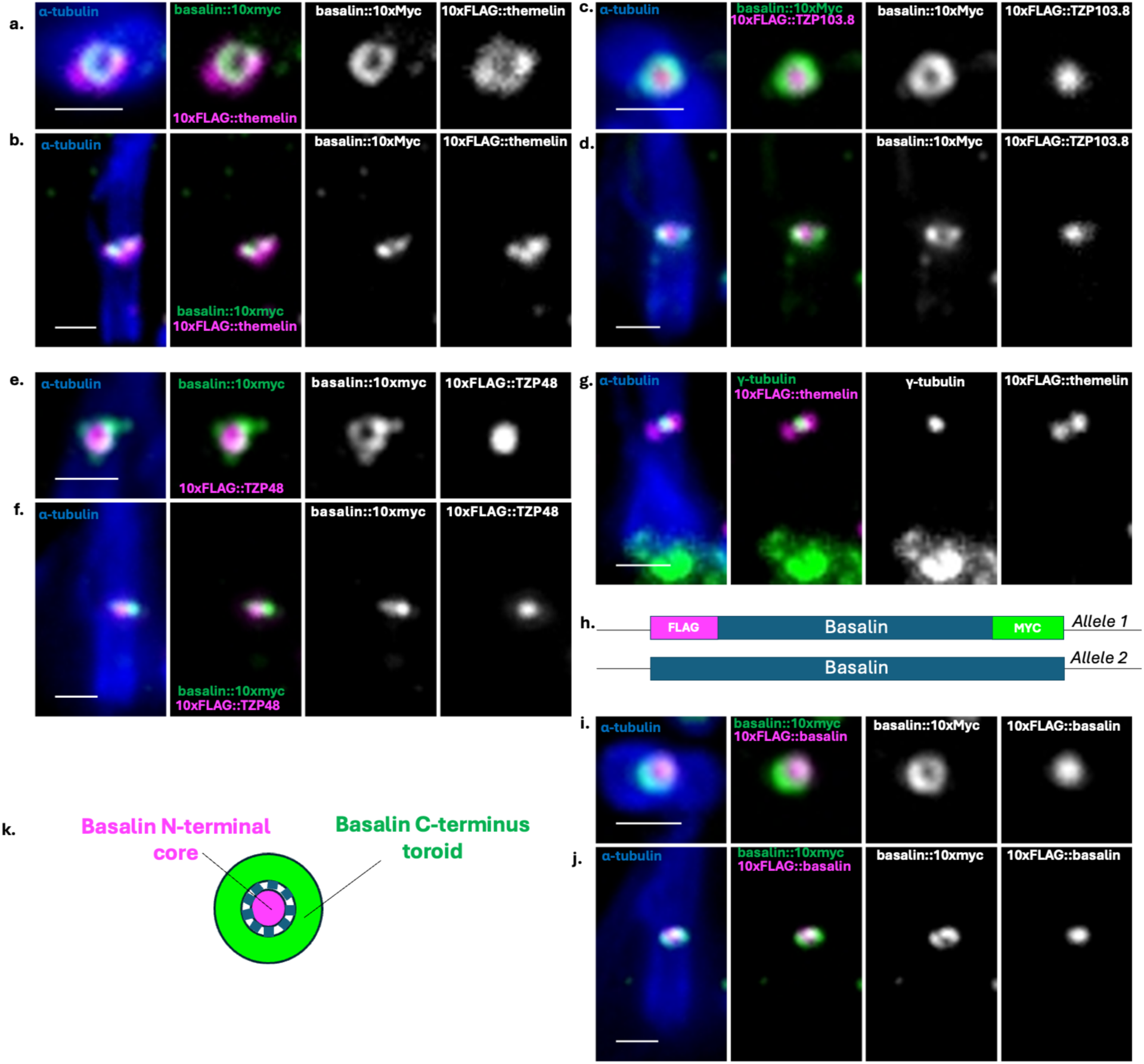
U-ExM reveals that CPAPs form nested rings surrounding a central core at the basal plate. Cells expressing CPAPs tagged with tandem epitopes (HA, myc or FLAG) were expanded and stained with anti-epitope antibodies to reveal their localisation at high resolution, and anti-tubulin as a structural reference. a, c, e, g, i) Basal plate cross sections; b, d, f, j) basal plate longitudinal sections. a, b) Basalin forms a ring nested within an outer themelin ring. In contrast, the basalin ring encircles a central core of TZP103.8 (c, d) and TZP48 (e, f). g) γ-tubulin localises to the base of the flagellum and the basal plate in the central core of the themelin ring. h-k) Bi-terminal tagging shows that the basalin C terminus forms a ring around an N terminal core. h) A cartoon showing a heterozygote where one allele of basalin is bi- terminally tagged with different epitopes. i, j) U-ExM analysis of cells co-expressing γ- tubulin::3xHA and 10FLAG::themelin reveals a small pool of gamma tubulin in the basal plate core in addition to the major pool at the base of the basal body. k) A model showing the ‘spoke-like’ arrangement of basalin at the basal plate. Scale bar = 1 μm.

The concentric arrangement of the CPAPs raised the question of whether individual proteins might span different compartments of the basal plate, with their termini oriented in distinct radial positions. To test this, we generated bi-terminally tagged cell lines for each CPAP, placing different epitope tags at the N- and C-terminus. Importantly, we confirmed by PCR that both tags were present on the same allele (Supplementary Fig. 2), ensuring that the signal from each tag reports the position of opposite ends of the same molecule rather than reflecting a mis-localisation artefact from tagging different alleles in *trans*. U-ExM of these lines revealed that for themelin, TZP103.8 and TZP48, both termini co-localised to the same compartment (Supplementary Fig. 3). Basalin, however, was the sole exception: its C terminus formed the ring, while its N terminus occupied the central core (Fig. 2g, h, j). Thus, basalin is radially oriented within the basal plate, bridging the outer ring and the inner core.

Although γ-tubulin has not been detected at the basal plate by standard fluorescence microscopy, the enhanced resolution afforded by U-ExM prompted us to revisit its localisation. U-ExM analysis of cells co-expressing γ-tubulin::3xHA and 10xFLAG::themelin resolved a small but distinct pool of γ-tubulin within the basal plate core, in addition to the major pool at the proximal end of the basal body (Fig. 2i). This basal plate pool had not been detected previously and provides the first direct evidence that γ-tubulin is physically present at the site of CP nucleation.

Together, these data reveal that the CPAPs are arranged as a set of nested rings - themelin outermost, basalin C terminus intermediate - surrounding an inner core composed of the basalin N terminus, TZP103.8, TZP48, and γ-tubulin.

### A subset of CPAPs is preassembled at the pBB

We next asked where the CPAPs are first recruited during the biogenesis of a new flagellum. In *T. brucei*, the probasal body (pBB) matures into the basal body that will nucleate the new flagellum and so represents the earliest structure at which CP assembly factors could be pre-positioned. To survey CPAP localisation, we tagged each protein with mNeonGreen and prepared detergent-extracted cytoskeletons for fluorescence microscopy. TZP103.8 appeared as a single spot at the flagellar base (Fig. 3b), consistent with its localisation exclusively at the mature basal plate. In contrast, basalin, themelin, TZP48 and γ-tubulin each appeared as two spots (Fig. 3a, c–e), raising the possibility that the second, fainter, spot corresponds to a pool at the pBB.

**Figure 3:**
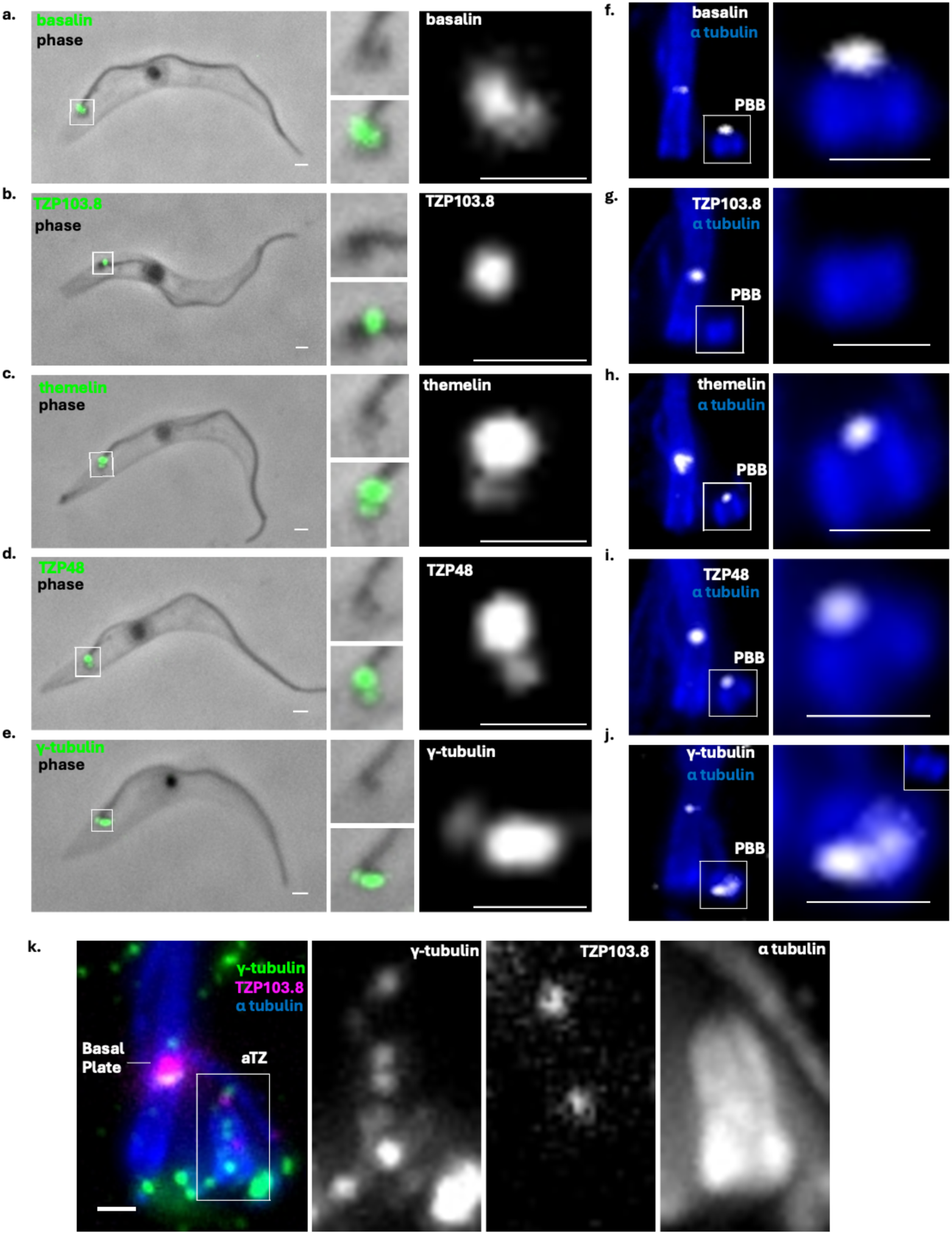
An immature central pair assembly complex resides on the distal pro- basal body. a-e) Basalin, TZP103.8, themelin, TZP48, and γ-tubulin were tagged with mNeonGreen and cells detergent extracted to prepare cytoskeletons for fluorescence microscopy. Insets highlight the signal at the base of the flagellum as either a single spot (TZP103.8) or two spots (basalin, themelin and TZP48, and γ-tubulin. f-j) Basalin, TZP103.8, themelin, TZP48, and γ- tubulin were tagged with tandem epitopes and cells detergent extracted to prepare cytoskeletons for U-ExM analysis. Insets show a distinct basalin, themelin and TZP48 signal at the distal end of the immature pro-basal body, but not for TZP103.8 or γ-tubulin. k) Cells co-expressing γ-tubulin::3xHA and 10xFLAG::TZP103.8 were analysed by U-ExM to assess the timing of TZP103.8 and γ-tubulin addition to the newly assembling TZ (aTZ). Scale bar = 1 μm.

To resolve the identity of these spots, we analysed tandem-epitope-tagged CPAP lines by U-ExM. This confirmed that basalin, themelin and TZP48 each localised to the distal end of the immature pBB somewhat offset to the microtubules, in addition to their established positions at the mature basal plate (Fig. 3f, h, i). By contrast, neither TZP103.8 nor γ-tubulin was detected at the pBB by U-ExM (Fig. 3g, k), indicating that these two proteins are recruited only later, during or after the pBB-to-BB transition. These data establish that a subset of CPAPs - basalin, themelin and TZP48 - form an immature CP assembly complex at the pBB before flagellum extension.

To define the timing of TZP103.8 and γ-tubulin addition more precisely, we co-expressed γ- tubulin::3xHA and 10xFLAG::TZP103.8 and examined cells at different stages of new TZ assembly by U-ExM. Both proteins appeared at the assembling TZ only once maturation was underway but before axoneme assembly (Fig. 3j). This places TZP103.8 and γ-tubulin recruitment downstream of the initial pBB complex, occurring during the final stages of basal plate assembly.

### iU-ExM resolves basalin and themelin into discrete puncta

To test whether the CPAP rings resolve into higher-order structures, we applied iU-ExM, which provides a further increase in effective resolution through successive rounds of expansion. iU-ExM of cells expressing 10xHA::themelin revealed that the themelin ring is not continuous, but in fact a lattice-like structure at the basal plate. Longitudinal sections showed this lattice in both tilted and perpendicular orientations relative to the axoneme (Fig. 4a, b), and different Z planes resolved the themelin signal into discrete puncta (Fig. 4c, d). iU-ExM of basalin::10xHA similarly resolved punctate substructure at the mature basal plate but with smaller diameter and height (Fig. 4e,f). Notably, punctate symmetry was evident for themelin and basalin at the pBB (Fig. 4d, e), demonstrating that this higher-order organisation is already established before the addition of TZP103.8 and γ-tubulin. Comparing the mature basal plates of old and new flagella in dividing cells revealed that Comparing the mature basal plates of old and new flagella in dividing cells revealed that the basalin signal, which is initially perpendicular to the long axis of the axoneme, becomes progressively more tilted as the basal plate ages (Fig. 4g).

**Figure 4.**
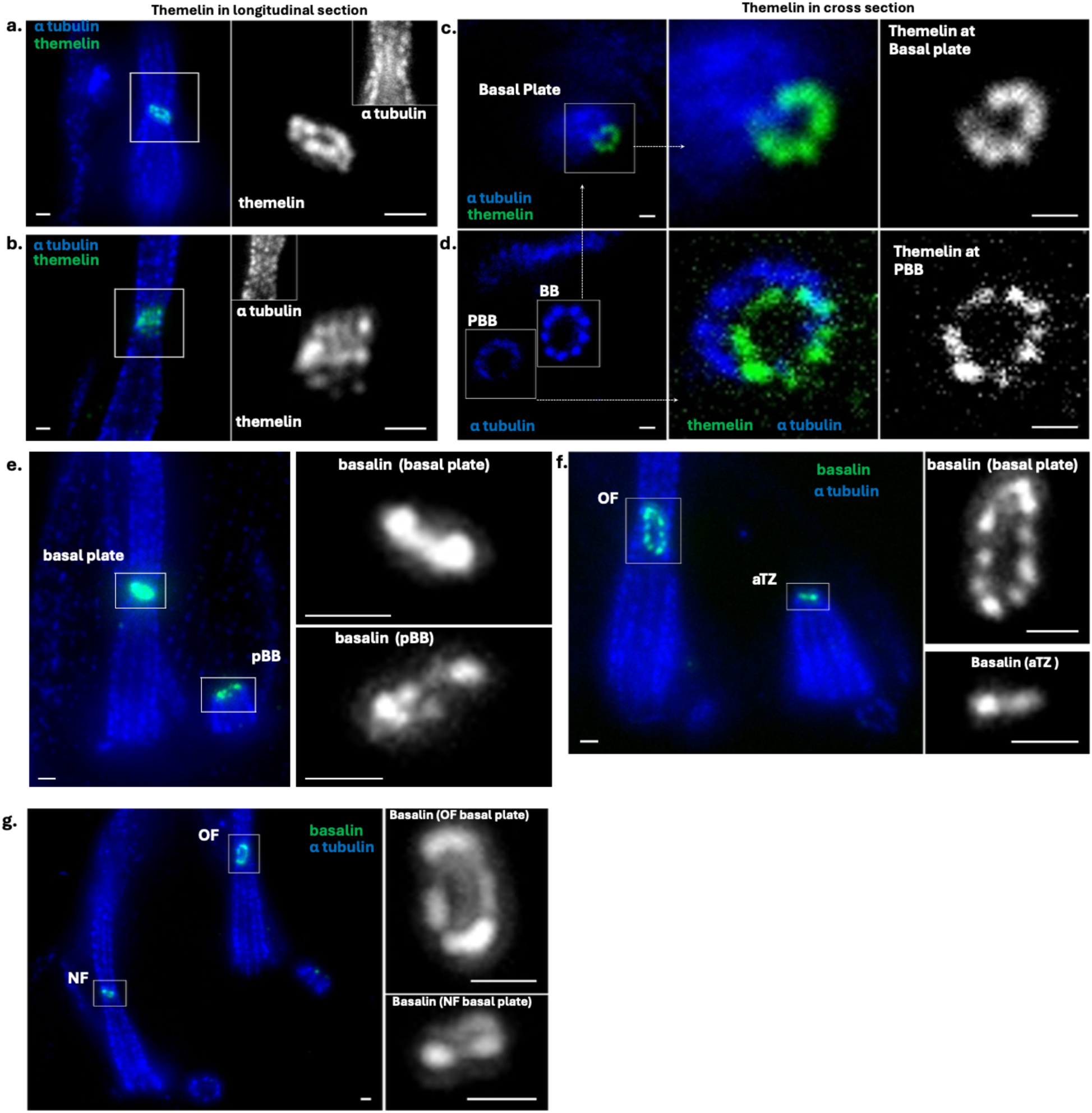
iU-ExM resolves basalin and themelin rings into punctate symmetry at the pBB and basal plate. Cells expressing 10xHA::themelin (a-d) and basalin::10HA (e-g) were analysed by iU-ExM. a and b) Longitudinal sections reveal themelin as a lattice-like structure at the flagellar basal plate. In a) the basal plate is tilted whereas in b) it is perpendicular to the axoneme. **c and d)** different Z planes reveal themelin as having punctate symmetry at the basal plate (c) and probasal body (d). e) Basalin signal exhibits discrete puncta or rings that is already established at the pBB. f) basalin signal at the basal plate of the old flagellum (OF) and an assembling new TZ (aTZ). g) Basalin signal at the basal plate of the old flagellum (OF) is more tilted than at the new flagella (NF) in dividing cells.

### Themelin is the master recruiter of an immature CP assembly complex on the distal probasal body

To establish the dependency relationships among the pBB-localised CPAPs, we generated cell lines co-expressing pairs of CPAPs with different tandem epitope tags and depleted each in turn by RNAi. Ablation of themelin had a profound effect on the other pBB-localised CPAPs: 35/39 pBBs lacking themelin had lost basalin, and 42/43 had TZP48 (Fig. 5e, f), indicating that themelin is required for the stable recruitment of both proteins. The converse relationship was notably weaker: ablation of basalin caused loss of themelin in fewer than half of affected pBBs (10/23; Fig. 5c, d), and a similar partial loss was observed for TZP48 (12/28 pBBs lacking basalin also lacked TZP48; Fig. 5a, b). Thus, while basalin contributes to the presence of both themelin and TZP48 at the pBB, it is not absolutely required, in contrast to its absolute requirement for TZP103.8 recruitment to the mature basal plate (16). Conversely, TZP48 depletion did not affect either themelin or basalin at the pBB (Fig. 5a, b). Together, these data establish themelin as the master recruiter of the immature CP assembly complex: its loss is sufficient to displace both basalin and TZP48 from the pBB, whereas basalin likely plays a contributory but subsidiary role in retaining themelin and TZP48 at this site.

**Figure 5:**
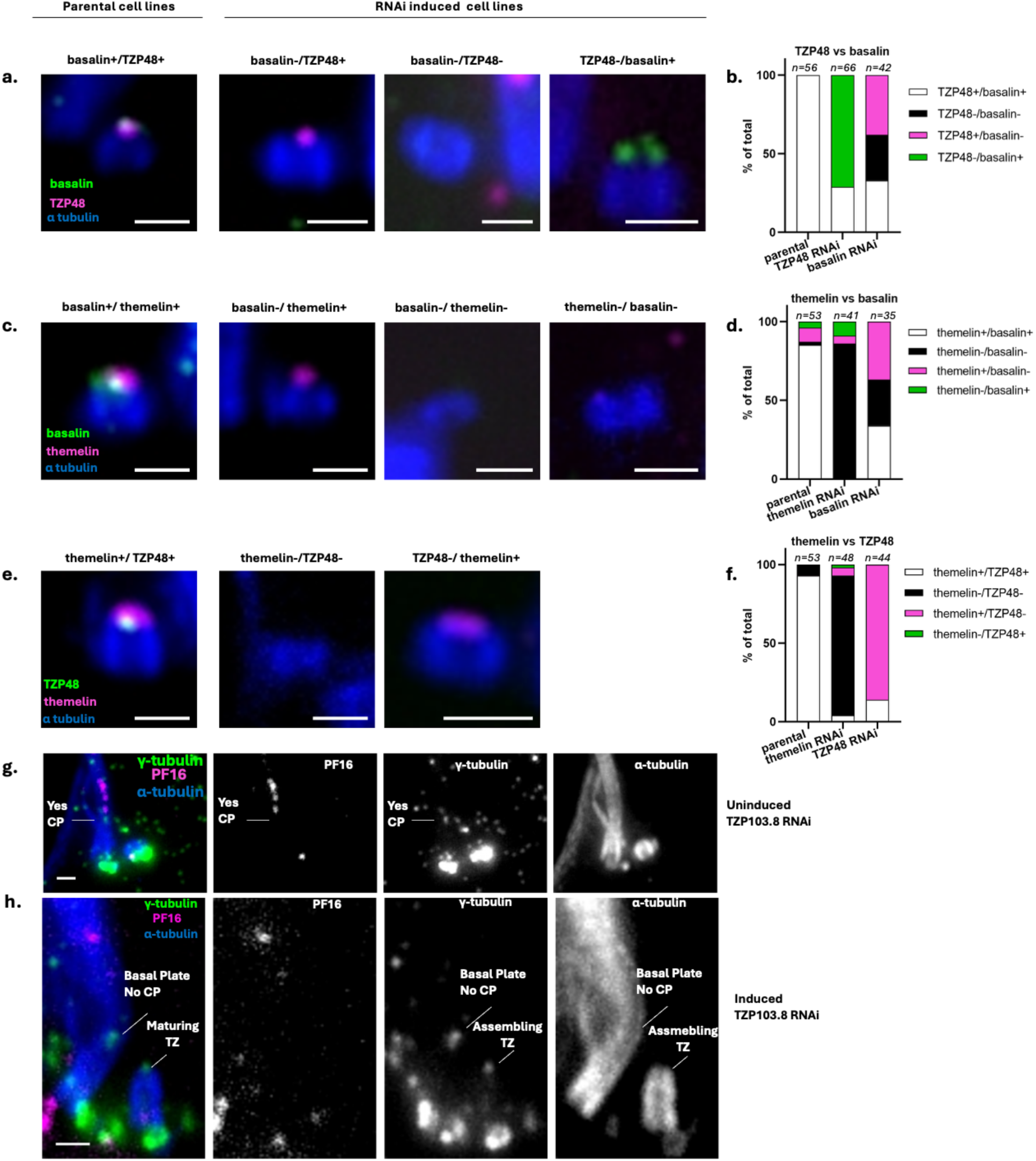
Themelin is the CPAP master recruiter at the pBB with TZ103.8 and gamma tubulin independently co-recruited during the final stages of basal plate maturation. Cells co-expressing two CPAPs tagged with different tandem epitopes were transfected with RNAi plasmids targeting one or other tagged CPAP and induced prior to U-ExM analysis. a-f) Examples (left panels) and quantification (right panels) of pro-basal bodies (pBBs) positive for both markers in control cells, or missing one or both markers in RNAi cells. a, b) Basalin and TZP48; c, d) basalin and themelin; e, f) themelin and TZP48. g, h) Cells co-expressing γ- tubulin::3xHA and PF16::3xFLAG were transfected with a TZP103.8 RNAi plasmid. g) A flagellum from uninduced cells showing a short pro-basal body and an intact central pair (PF16-positive). h) In dividing TZP103.8 knockdown cells, γ-tubulin signal is retained at the basal plate of mature flagella that lack the central pair, and the distal end of the assembling transition zone. Scale bar = 1 μm.

We previously showed that ablation of γ-tubulin does not prevent TZP103.8 recruitment to the mature basal plate (16). To test the converse, we knocked down TZP103.8 in cells co-expressing γ-tubulin::3xHA and PF16::3xFLAG and examined them by U-ExM. Loss of TZP103.8 from the basal plate was confirmed by the absence of axonemal PF16 signal, indicating that the CP had not formed. Nevertheless, γ-tubulin was retained at both the basal plate and the distal end of the nascent TZ (Fig. 5g, h), demonstrating that γ-tubulin alone is not sufficient for CP nucleation and that TZP103.8 functions as a critical downstream effector.

### Themelin ablation causes redistribution of CPAPs into ectopic supernumerary hubs

Predicting that themelin loss would also disrupt the localisation of CPAPs at the mature basal plate, we ablated themelin by RNAi in cell lines co-expressing pairs of fluorescently tagged CPAPs and examined the consequences by fluorescence microscopy. In uninduced control cells, basalin and TZP103.8 co-localised as expected at the flagellar base, with basalin additionally visible at the pBB (Fig. 6a, c). Following 72 hours of themelin RNAi induction, the pBB basalin and TZP48 signal was lost and, unexpectedly, co-tagged basalin and TZP48 colocalised into multiple ectopic puncta along the length of the flagellum (Fig. 6b, d). The same redistribution was observed for basalin and TZP103.8 (Fig. 6c, d), indicating these represent supernumerary hubs containing all three CPAPs. Reproducing this experiment with fluorescently tagged basalin and PF16 confirmed that flagella containing supernumerary CPAP hubs lacked axonemal PF16 signal, demonstrating that these ectopic assemblies are non-functional and do not nucleate CPs at aberrant positions (Fig. 6e, f). Quantification showed that supernumerary basalin hubs accumulated progressively in assembling flagella over 72 hours of themelin RNAi depletion (Fig. 6g).

**Figure 6.**
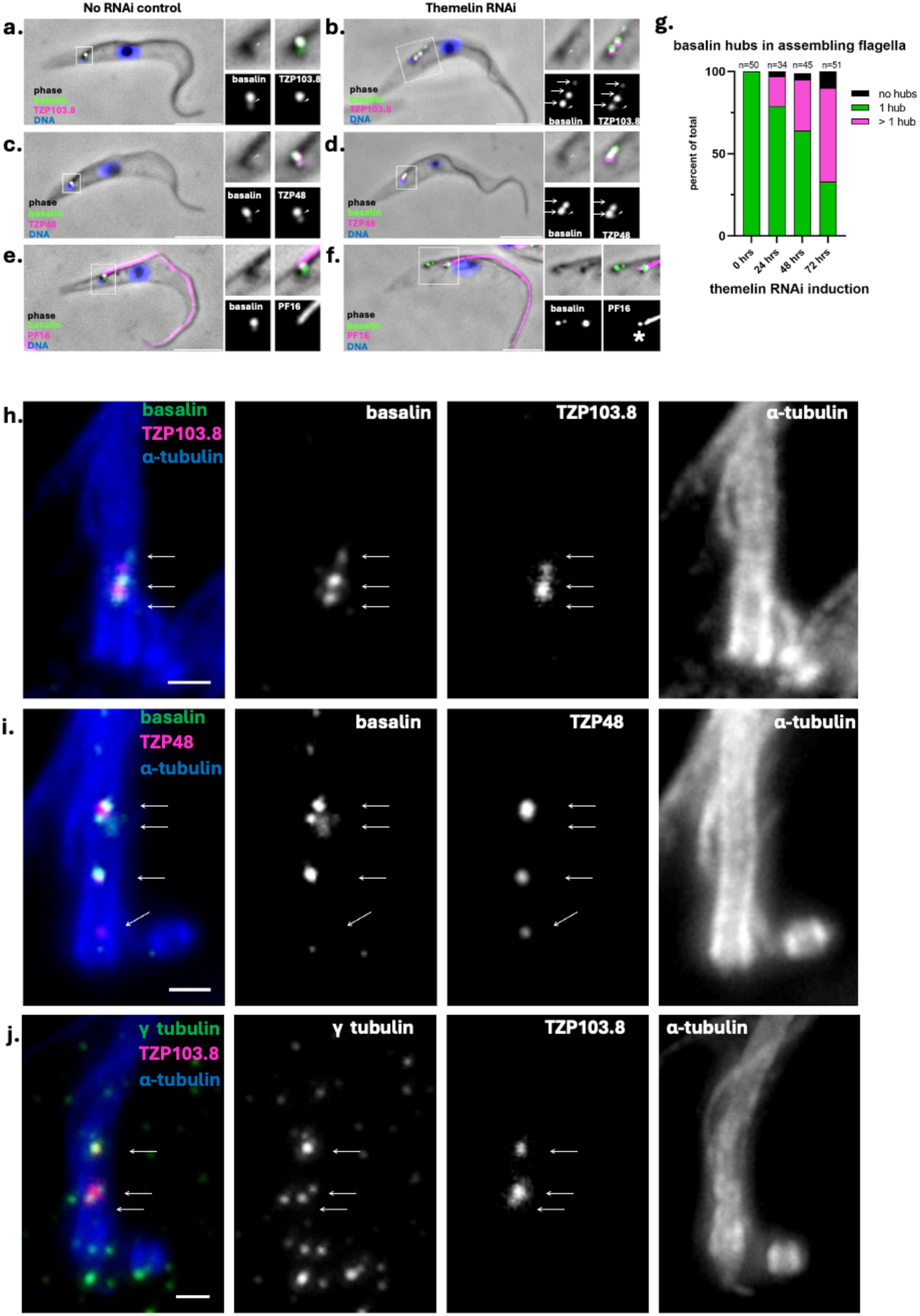
Themelin RNAi ablation causes redistribution of CPAPs into ectopic supernumerary hubs inside the flagellum. Cells co-expressing CPAPs tagged with either fluorescent proteins (a-g) or tandem epitope tags (g-i) were induced for themelin RNAi for 72 hours and analysed by either fluorescence microscopy or U-ExM, respectively. a-b) basalin::mNeonGreen and mScarlet-I::TZP103.8. c-d) basalin::mNeonGreen and mScarlet-I::TZP48. e-f) basalin::mNeonGreen and PF16::mScarlet-I. h) basalin::10xMyc and 10xFLAG::TZP103.8. i) basalin::10xMyc and 10xFLAG::TZP48. j) γ-tubulin::3xHA and 10xFLAG::TZP103.8. Arrowheads indicate the pBB signal (a, c) or signal absence (b, d). g) Quantification of basalin hubs in control and RNAi cells based on fluorescence microscopy data. Arrows indicate supernumerary hubs. Scale bars: a-f = 5 μm, g-i = 1 μm.

To characterise the ultrastructure of these ectopic puncta, we examined themelin-ablated cells by U-ExM. U-ExM of co-tagged cells revealed that the ectopic puncta corresponded to either discrete hubs or chains of CPAP aggregates, within the axoneme in which basalin, containing TZP103.8, TZP48 and γ-tubulin (Fig. 6h–j). However, within these accumulations, basalin did not form its characteristic ring encircling TZP103.8 and TZP48 seen at the native basal plate. Thus, in the absence of themelin, although the remaining CPAPs retain an intrinsic capacity to co-assemble and be imported into the flagellum, key features of their spatial arrangement are lost. These data indicate that themelin is required both for targeting the CP assembly complex to its correct location and for organising the concentric architecture of the CPAPs within it.

### Anterograde IFT is required for efficient incorporation of TZP103.8, γ-tubulin and basalin into the basal plate

IFT172 knockdown blocks axoneme assembly, producing axoneme-less TZ ’stubs’ that nevertheless retain a basal plate by EM. However, whether the basal plate CPAPs are correctly incorporated is unknown. Given that the pBB complex carries substantially less CPAP signal than the mature basal plate (Fig. 3a-j), and that the addition of TZP103.8 and γ- tubulin represents the final step that triggers CP nucleation (Fig. 3k), we asked whether anterograde IFT provides a mechanism for accumulating CPAPs to functional levels and delivering the remaining components.

To assess this, we knocked down a key component of anterograde IFT, IFT172, in cells co-expressing epitope-tagged TZP103.8 and γ-tubulin for U-ExM analysis (Fig. 7a-c; Supplemental Fig. 3). As expected, all non-induced cell TZs were strongly positive for TZP103.8 (39/39) and either strongly positive (31/39), or faintly positive (8/39) for γ-tubulin. In contrast, analysis of TZ stubs after 72 hours IFT172 RNAi, the first timepoint at stubs are discernible, revealed that the majority had either no detectable or only faint signal for both markers. For γ-tubulin, 33/56 (58%) of stubs were negative and a further 17/56 (30%) were only faintly positive, with just 6/56 (10%) retaining clear signal. TZP103.8 showed a similar distribution, with 22/56 (38%) of stubs negative and 23/56 (41%) faintly positive, leaving only 12/56 (21%) with detectable signal.

**Figure 7.**
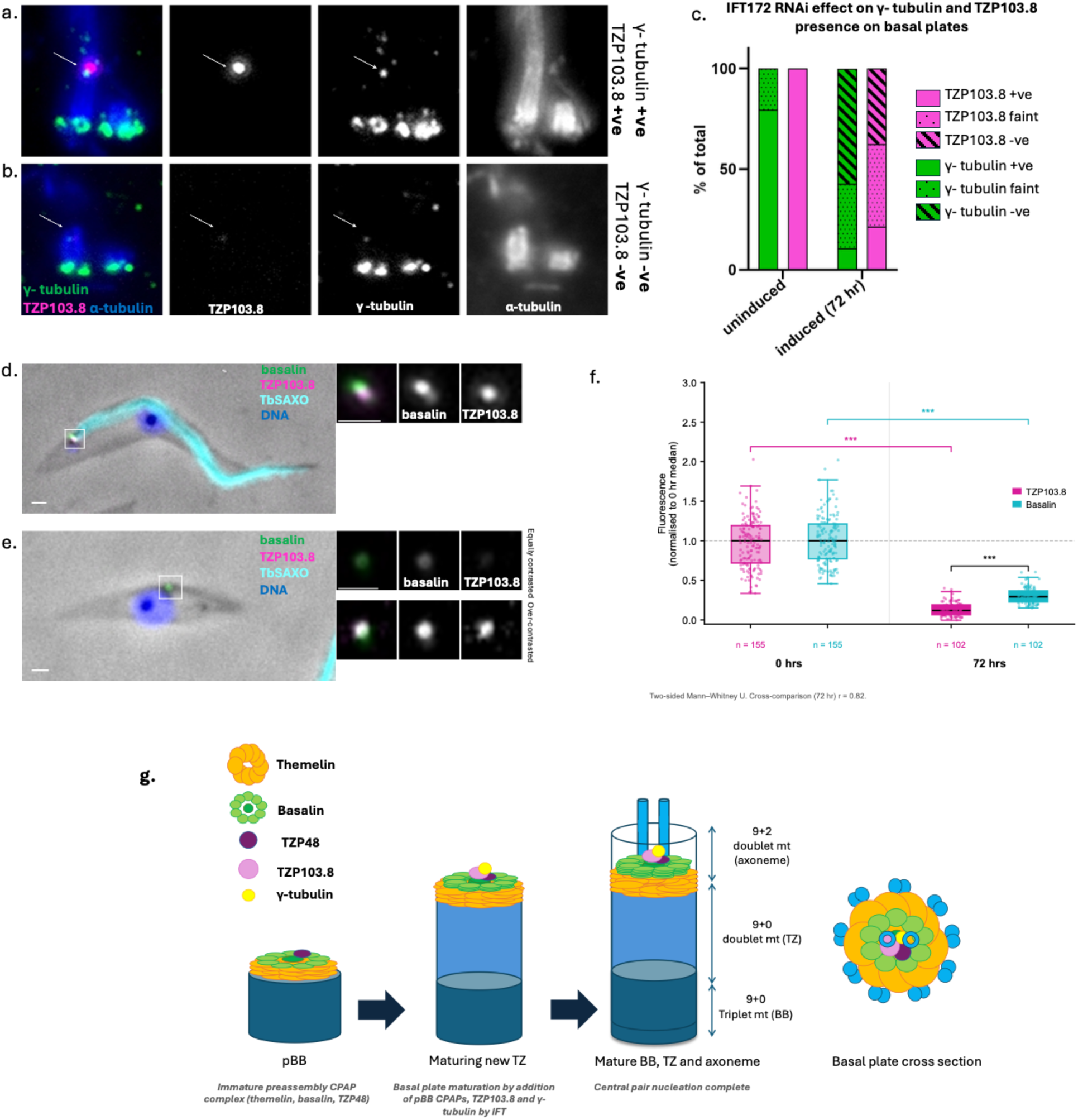
IFT is required for efficient transport of CPAPs and γ-tubulin to the basal Plate. Cells expressing tagged CPAPs and γ-tubulin were induced for IFT172 RNAi for 72 hours and analysed for basal plate localisation. a-c) U-ExM analysis of gamma tubulin and TZP103.8 at TZs after IFT172 RNAi. Representative U-ExM of flagellar TZs from controls (a) and axoneme- less TZ stubs from IFT172-depleted cells (b) expressing γ-tubulin::3xHA and 10xFLAG::TZP103.8. c) Quantification reveals that, in contrast to controls, many TZ stubs have undetectable, or barely detectable, levels of basalin and gamma tubulin after 72 hours IFT172 ablation (underlying data shown in Supplementary Fig. 4). d-f) Fluorescence microscopy analysis of TZP103.8 and basalin at TZs after IFT172 depletion. Uninduced (d) and IFT172 RNAi-induced (e) cells co-expressing basalin::mNeonGreen and mScarlet- I::TZP103.8, were stained with anti-TbSAXO to identify cells lacking an axoneme in induced cultures. Insets show regions of interest at higher magnification; upper insets in h are shown at the same contrast settings as (d) to highlight lower protein levels, lower insets are shown at high contrast. f) Fluorescence intensity of basalin::mNeonGreen and mScarlet- I::TZP103.8 measured at flagellar TZs from control cells, and axoneme-less stubs from IFT172-depleted cells, normalised to the median of the respective control condition after background subtraction (Supplemental Data 2). Boxes show the median (central line) and interquartile range (IQR, 25th–75th percentile); whiskers extend to the most extreme data point within 1.5 × IQR of the box. All individual data points are overlaid (jittered horizontally for clarity). Data from one representative experiment is shown (n=2). Statistical comparisons used the two-sided Mann–Whitney U test: 0 hrs vs 72 hrs for each protein, and TZP103.8 vs basalin at 72 hrs. *** p < 0.001. The effect of knockdown was significantly greater on TZP103.8 than on basalin (rank-biserial effect size r = 0.82). Scale bars: a-d = 1 μm, g-h = 5 μm. g) A model of how the central pair assembly complex is formed.

To assess whether IFT-dependent basal plate maturation extends to pBB-preassembled CPAPs, we reproduced the IFT172 RNAi in cells co-expressing fluorescently tagged TZP103.8 and basalin, and quantified fluorescence signal intensity at TZ stubs identified by the absence of the axoneme marker TbSAXO (Fig. 7d-f). TZP103.8 signal intensity was significantly reduced following IFT172 RNAi (88%), yet remained detectable in most TZ stubs, suggesting that the greater sensitivity of widefield fluorescence microscopy permits detection of a residual IFT-independent pool that falls below the detection threshold of U-ExM. Basalin signal was also reduced (77%), suggesting that even CPAPs first recruited at the pBB stage are subject to IFT-dependent accumulation during basal plate maturation.

## Discussion

The central pair of microtubules is essential for flagellar motility, yet the molecular events that lead to its nucleation have remained largely obscure. In this study, we identify two new central pair assembly proteins - themelin and TZP48 - and define the spatial organisation, temporal hierarchy and delivery mechanisms of the CPAPs and γ-tubulin in *T. brucei*. Our findings reveal that preparation for CP assembly initiates at the probasal body with the formation of a pre-assembly complex that subsequently matures through the staged addition of further components.

A central finding of this work is the identification of themelin as the master recruiter of the pBB complex. Themelin is required for the recruitment of both basalin and TZP48 to the pBB, whereas basalin makes only a partial contribution to the retention of the other pBB-localised CPAPs. Themelin is also required for organising the concentric architecture of the CPAPs, as its loss produces supernumerary hubs in which basalin, TZP103.8, TZP48 and γ-tubulin co-assemble but without the correct spatial arrangement. This dual role - in both positioning and organising the complex - distinguishes themelin from a simple recruitment factor and suggests it acts as an architectural template.

U-ExM and iU-ExM reveal that the basal plate possesses a degree of internal order that was not apparent by conventional microscopy or EM (17, 22). The CPAPs are arranged as nested rings surrounding a central core containing the key, downstream effector proteins. Interestingly, basalin connects the core compartment to the ring, with its C terminus in the ring and N terminus in the core; the path taken between these anchor points is unresolved, and may be radial, oblique, or - given the predicted disorder of the intervening sequence - conformationally heterogeneous. iU-ExM further resolves the themelin and basalin rings into punctate arrays with radial symmetry, with themelin forming a lattice-like structure. This punctate organisation is already present at the pBB, indicating that the symmetry of the complex is established early and is not imposed by later events such as TZ maturation or CP nucleation. The observation that the basalin signal becomes progressively more tilted in older basal plates is intriguing and may reflect continued structural remodelling of the basal plate over time, perhaps in response to mechanical forces exerted by the growing or beating axoneme.

The biochemical underpinning of how two singlet CP microtubules are nucleated at the basal plate remains an open question. γ-tubulin is intrinsically a poor microtubule nucleator. In other systems, GCP2 and GCP3 associate with two γ-tubulins to form the heteromeric γ-tubulin small complex (γTuSC), and multiple γTuSCs are then oligomerised into the cone-shaped γ-tubulin ring complex (γTuRC), in which GCP4, GCP5 and GCP6 occupy specific positions within the spiral that are essential for its asymmetric architecture (23). CM1 domain-containing proteins such as CDK5RAP2 then promote conformational closure of the γTuRC to match the 13-protofilament geometry of the microtubule (24, 25). In organisms that lack GCP4–6, such as budding yeast, the CM1 motif of Spc110 instead drives γTuSC oligomerisation into a ring at the spindle pole body (24). Trypanosomes encode GCP2, 3 and 4, but lack detectable orthologues of GCP5 and GCP6 and CM1 domain-containing proteins, and GCP4 is dispensable for CP nucleation (21), indicating that trypanosomes can neither assemble a canonical γTuRC nor use CM1-driven oligomerisation to form one. Given that our fluorescence microscopy shows that TZP103.8 is in substantial excess of γ-tubulin, it is unlikely that TZP103.8 is a non-canonical γTuRC component. In this context, our finding that γ-tubulin localises to the basal plate in TZP103.8-depleted cells yet fails to nucleate the CP has important implications for models of CP nucleation. We propose instead that TZP103.8, as the final CPAP recruited to the basal plate, provides the structural environment that constrains γTuSC into the precise geometry required to nucleate two microtubules in the correct positions and orientations for central pair formation. Additionally, TZP103.8 may also stabilise nascent microtubules during the critical early seconds of nucleation, when only a few tubulin dimers have been added and the assembly is vulnerable to collapse. Whether nucleation occurs from individual, non-oligomerised γTuSCs positioned within the basal plate structural environment, or requires an as-yet-undiscovered mechanism of γTuSC oligomerisation, remains to be determined.

Our data reveal that anterograde IFT is required for efficient accumulation of CPAPs and γ- tubulin at the basal plate, aligning with TEM data showing that the TZ and basal plate are structurally reduced in IFT172 and IFT88 knockdown cells (26, 27). Notably, this IFT- dependency extends beyond the late-arriving proteins to include basalin, a pBB- preassembled CPAP, indicating that IFT contributes to overall basal plate maturation rather than solely delivering the final components. This suggests a role for IFT in assembling the distal TZ both by delivering CPAPs and by promoting extension of the transition zone microtubules on which they are assembled.

Although reduced, residual CPAP signal nonetheless persists at TZ stubs where IFT is clearly insufficient to support axoneme assembly, suggesting that IFT functions to accumulate CPAPs beyond the levels provided by pBB assembly and diffusion alone. Given that our iU-ExM data show that the punctate 9-fold architecture of the complex is already established at the pBB, IFT does not remodel the structure but loads additional protein onto a pre-existing template, progressively building the structural platform upon which TZP103.8 and γ-tubulin are assembled into the geometric arrangement required for CP nucleation. More broadly, these data implicate IFT in a non-axonemal process at the TZ, consistent with evidence that IFT contributes to TZ maintenance in mammalian RPE1 cells (28). Together, these observations suggest that IFT plays a broader role in TZ biogenesis and maturation than has been generally appreciated.

Basal plate-like structures are widespread at the base of motile cilia (29), yet how these structures organise γ-tubulin for CP nucleation remains poorly understood in any system. The assembly logic described here - early templating at a precursor structure followed by staged recruitment of structural and nucleator components – may therefore represent a general principle for how motile cilia solve the problem of CP nucleation. More broadly, the strategy of pre-assembling a template complex at a precursor organelle, then licensing its activity through the staged addition of effector components, may be a recurring mechanism for building complex molecular machines.

## Material and Methods

### Cell lines generation and cultures

*Trypanosoma brucei* TREU927 SMOX P927 cells (30) and TREU927 cells genetically modified using pJ1339 (31) were grown at 28^°^C in SDM79 (32). Genetically modified knockout and protein tagged trypanosome cell lines were generated as described (33–35). RNAi was induced using 5-10 μg.mL^−1^ doxycycline using gene fragments cloned into pQuadra (36). Genomic DNA for PCR verification of gene deletion and biterminal tagging was prepared following manufacturer instructions (Mouse Direct PCR Kit, SELLEK, UK) using gene-specific RNAi primers (18) or primers designed to anneal to sequence encoding each epitope tag.

### Immunofluorescence and U-ExM

For immunofluorescence, 0.5% NP-40 in PEME treated ‘cytoskeletons’ were prepared, fixed in methanol and rehydrated in phosphate-buffered saline (PBS) as described (37). Slides were then incubated in block (2% fetal calf serum, 0.2 % TWEEN, in PBS) for 30 min, stained with primary antibodies or imaged immediately for fluorescent protein tags, prior to mounting in phosphate-buffered glycerol and imaging. Images were acquired as described in Paterou et al., 2025 using a Leica DM6 B microscope equipped with a metal halide lamp (EL 6000, 11504115) serving as the illumination source, a 63× objective with a numerical aperture of 1.30 (11506385) and a Leica K5 Microscope Camera (11547112).

U-ExM was performed as described (38–41). In brief, anchoring was performed either in whole cells or cytoskeletons using 0.7% electron microscopy grade formaldehyde (FA) and 1% acrylamide in PBS, at room temperature overnight. All prepared gels were stained with antibodies by immunofluorescence in 2% BSA/0.2% Tween in PBS overnight and washed with double distilled water and PBS. Primary antibodies used were: rabbit anti-HA (Cell signalling, clone C29F4), rabbit anti-Myc (Absolute Antibodies, clone 9E10), mouse monoclonal anti-FLAG (Sigma, clone M2), guinea pig anti-alpha-tubulin (ABCD antibodies, ABCD_AA345), secondary antibodies used were: anti-guineapig Alexa-555, anti-rabbit Alexa-488, anti-rabbit Alexa-647, anti-mouse Alexa-647, anti-mouse Alexa-488 (Invitrogen). Immunostained gels were imaged with cells facing down on poly-L-lysine (Sigma) treated glass-bottomed confocal dishes (Avator) at a widefield Leica inverted microscope DMi8 using a 100x oil lense, a metal halide light source and ORCA-flash 4.0 sCMOS camera (HAMAMATSU).

### Iterative U-ExM (iU-ExM)

Iterative ultra expansion microscopy (iU-ExM) was performed as described (42), with minor modifications. In brief, 4 million procyclic *T. brucei* cells were washed twice in vPBS (8 g/L NaCl, 0.22 g/L KCl, 2.27 g/L Na2HPO4, 0.41 g/L KH2PO4, 15.7 g/L sucrose, 1.8 g/L glucose, pH 7.4) and attached to a clean round coverslip (12 mm) before anchoring with 1.4% electron microscopy grade (FA) and 2% acrylamide (AA)in PBS, in the dark at room temperature overnight. The coverslip was then placed in the centre of an ice cold humidity chamber for gelation with monomer solution 1 (MS1) (10% AA, 19% Sodium Acrylate (SA, AK Scientific), 0.1% DHEBA (SIGMA), 0.25% tetramethylethylenediamine (TEMED, SIGMA), 0.1% Ammonium Persulfate (APS, SIGMA)) added to the top of the 12 mm coverslip with the cells, left for 15 minutes on ice, and then moved to 37°C in a humidified incubator for 45 min to polymerise. After gelation, the gels were denatured in denaturation buffer (200 mM SDS, 200 mM NaCl, 50 mM Tris-base, pH 6.8) in a 1.5 mL microfuge tube using a heat block at exactly 85°C for 90 min. After denaturation the gels were expanded by washing in double distilled water several times until gels reach a plateau of expansion (usually 5-fold expansion of the original size), then cut into 2 cm × 2 cm squares, and stained with antibodies (concentrations were double that used for U-ExM) shaking at 37°C at room temperature, before proceeding to the second expansion step. Gels were then washed in ice cold activated neutral gel solution (MS2) (10% AA, 0.05% DHEBA, 0.1% APS/TEMED in ddH2O) and incubated in a humidity chamber for 1 h at 37°C. Then the gels were anchored with 1.4% FA/2% AA in PBS shaking at 37°C overnight. Gels were then washed in 1× PBS for 30 min (3 × 10 min washes). The second gelation was performed in a 12 well plate with 1.5 mL of MS3 (10% AA, 19% SA, 0.1% BIS (SIGMA), 0.1% APS/TEMED), changing the solution 4- 5 times for a total of 30 min while shaking on ice with open lid to slow down polymerisation of the acrylamide. The gel was then moved on top of a glass slide, removing the excess solution with paper tissues, covered with a 22 × 22 mm coverslip and incubated in a humidity chamber for 1 h at 37°C for final polymerization. The entire gel was then incubated in 200 mM NaOH solution for 1 h in a small beaker shaking at RT, followed by washes with PBS for 30 min (3 × 10 min washes) until the pH dropped to pH 7 (checked with a pH strip). The gels were then washed several times in ddH2O and left in water in the cold overnight before imaging. Imaging was performed as described for U-ExM.

### Image analysis

Images were imported into OMERO (43) and analysed either in OMERO.figure, OMERO.web, or Fiji (44). Fluorescence intensity of spots (Fig. 6) was measured using a custom script (Supplemental data 2).

## Acknowledgements

AP and SD performed the experiments, analysed the data and co-wrote the manuscript. SD conceived and administered the project, provided supervision and funding. AP was supported by an Academy of Medical Sciences Springboard award (SBF006/1126) and BBSRC NIRG (BB/Y012739/1) to SD. We thank Marie Zelená and Dr Vladimír Varga (ICMG Czech Academy of Sciences) for their help establishing U-ExM, and Dr Philippe Bastin and Dr Aline Araujo Alves (Institut Pasteur) for their help establishing iU-ExM. We thank Professor Anne Straube (Warwick) for helpful comments throughout the project, Professor Keith Gull (Oxford) for critical reading of the manuscript, and WMS for their departmental support. The Warwick Advanced Bioimaging RTP facility provided help with sample preparation for TEM.

## Data Availability

All data supporting the findings of this study are available within the manuscript and its supplementary materials.

---

**Supplemental Figure 1.**
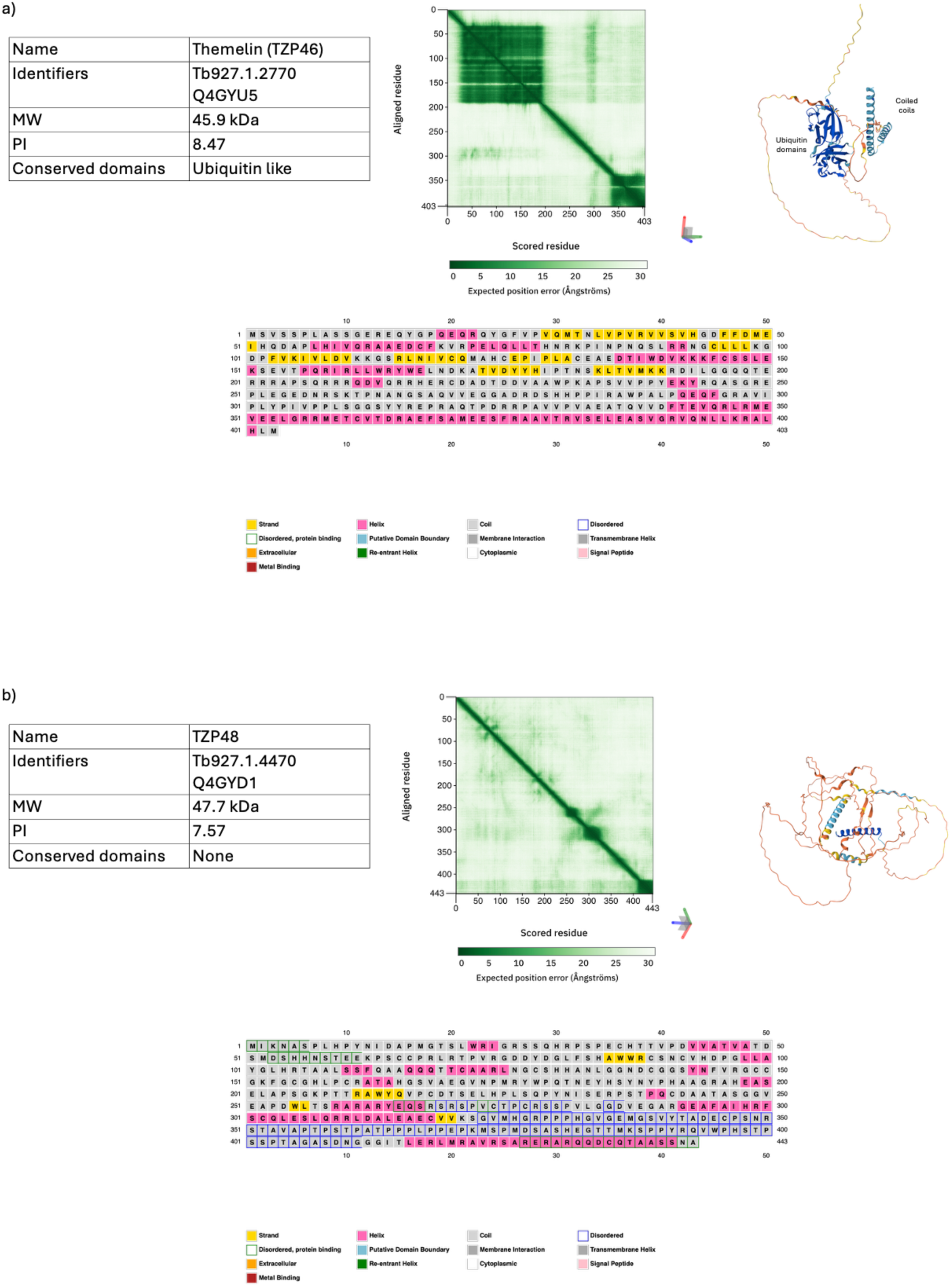
Predicted biochemical attributes of themelin and TZP48. Calculated biochemical attributes and structural predictions of themelin (a) and TZP48 (b) (45, 46).

**Supplemental Figure 2.**
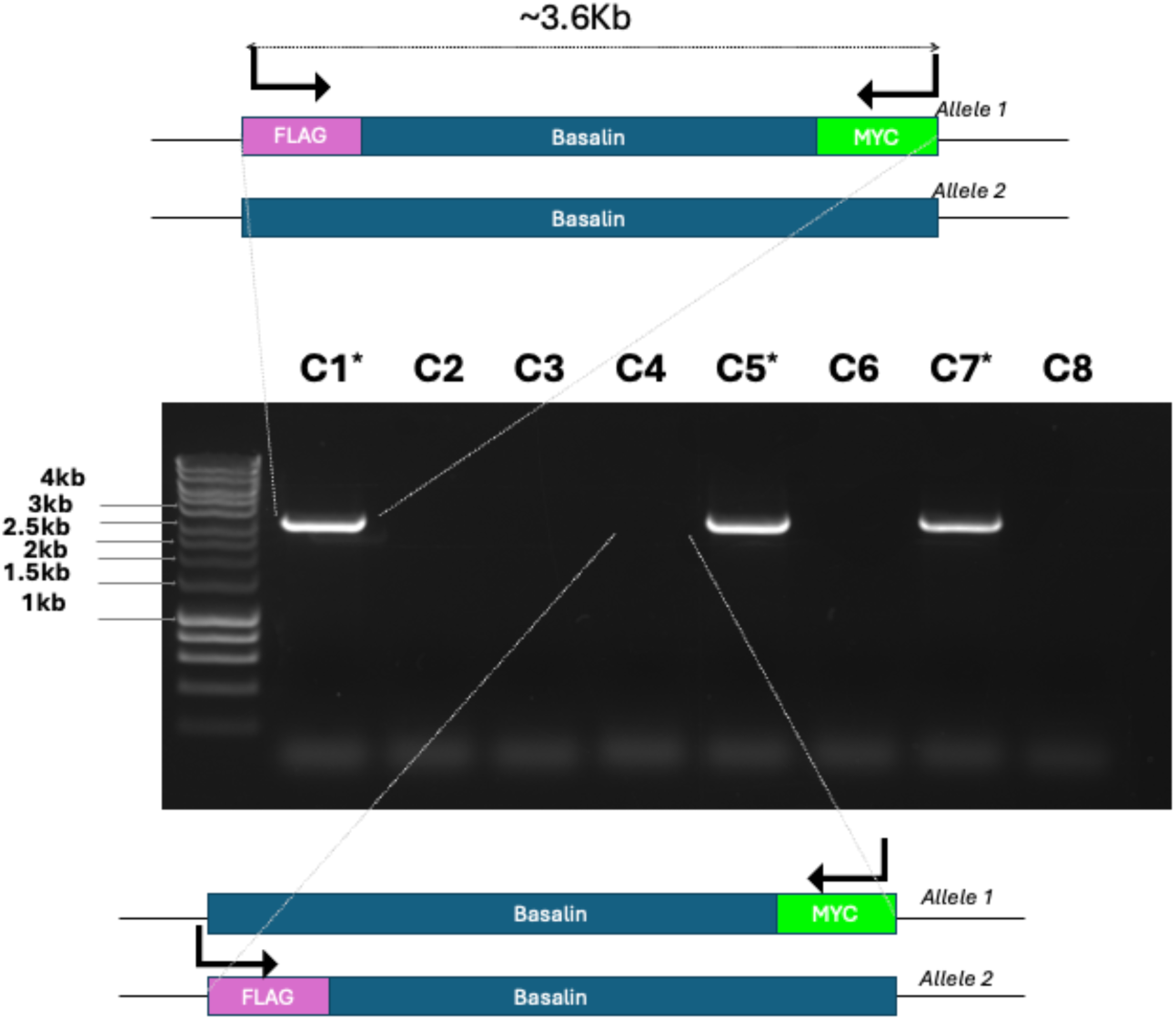
Screening for biterminally tagged basalin alleles. A cell line expressing basalin tagged with 10 tandem copies of the myc epitope tag at the C-terminus (basalin::10myc) was transfected to tag the basalin N-terminus with ten FLAG epitope tag (10FLAG::basalin). Genomic DNA prepared from drug resistant clones was used in a PCR with primers complementary to DNA encoding each epitope tag. Hence, only clones that have the same allele tagged with both the N-terminal FLAG and the C terminal myc (i.e. biterminal cis tagging) give rise to a PCR product. In contrast, if different alleles are tagged (biterminal trans-tagging) PCR does not occur. Here, of eight resistant clones screened, three clones gave rise to a PCR product (noted with an asterisk); clone 1 (C1) was sequenced to confirm successful biterminal cis-tagging and protein localisation verified by iimmunofluorescence (data not shown), and then taken forward for U-ExM analysis.

**Supplemental Figure 3.**
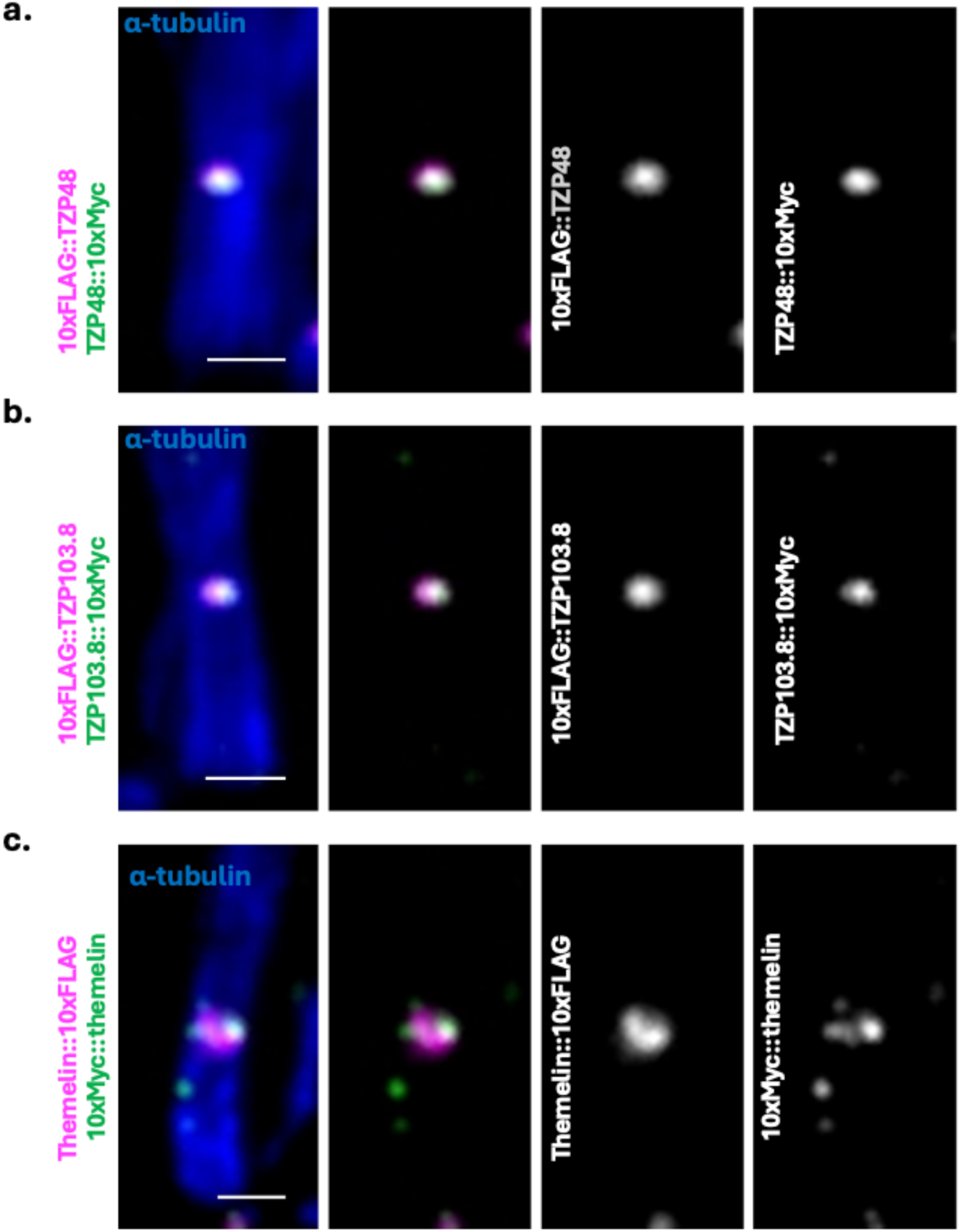
Double termini tagged themelin, TZP48 and TZP103.8 show similar pattern. U-ExM microscopy on cells expressing biterminally cis-tagged a) TZP48, b) TZP103.8 and c) themelin. All three CPAPs showed the same localisation whether they were tagged at the N-terminus or the C-terminus.

**Supplemental Figure 4.**
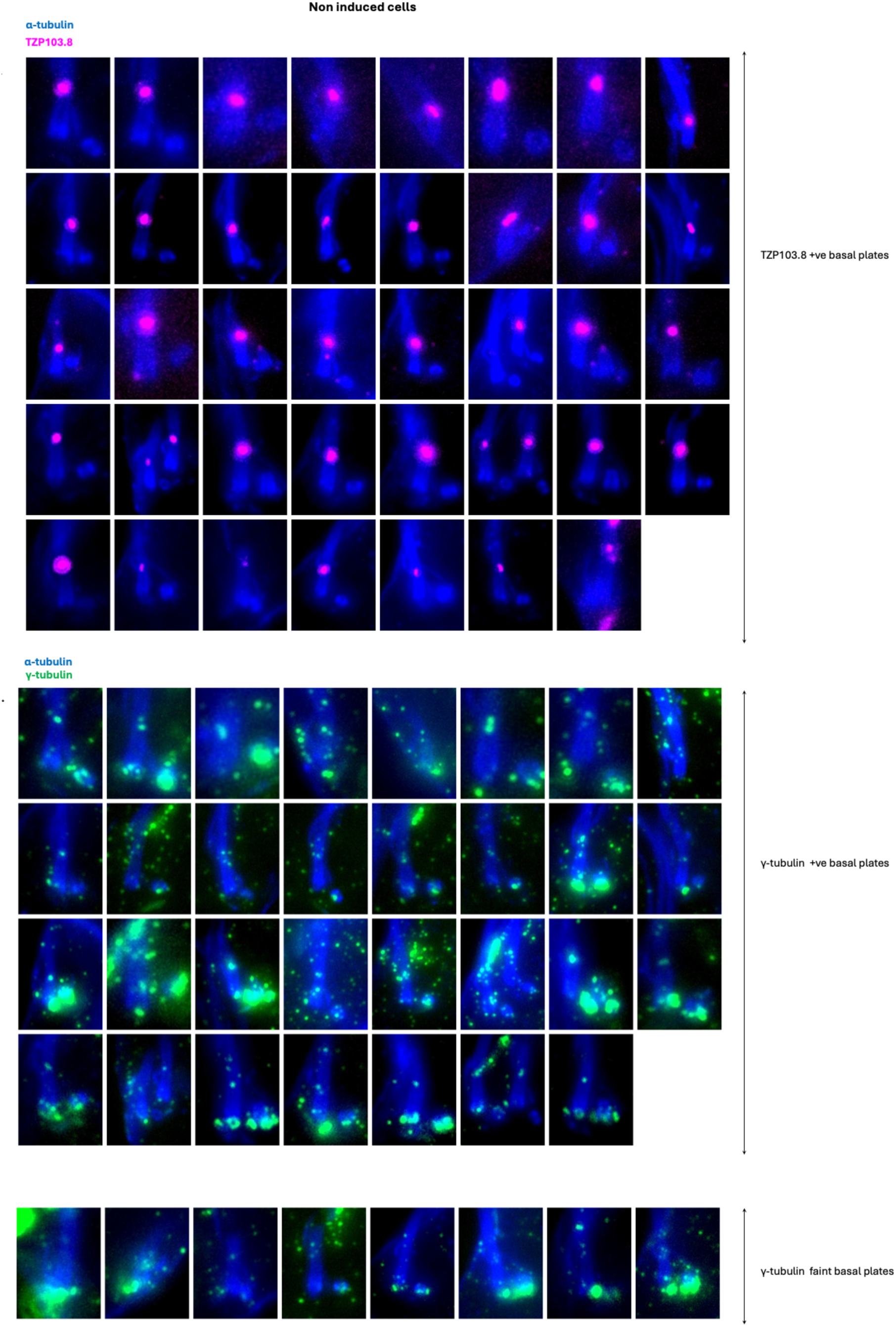
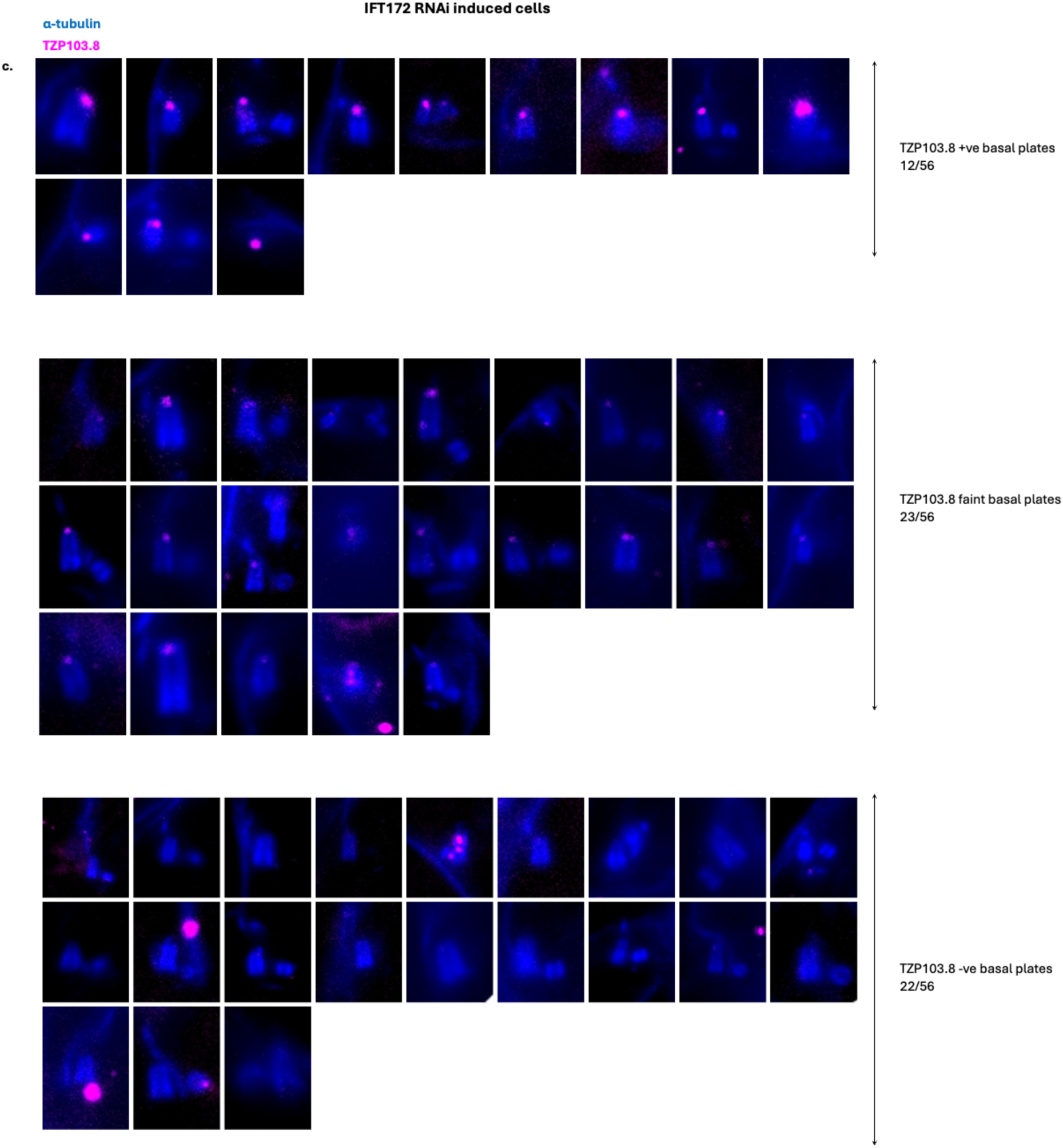
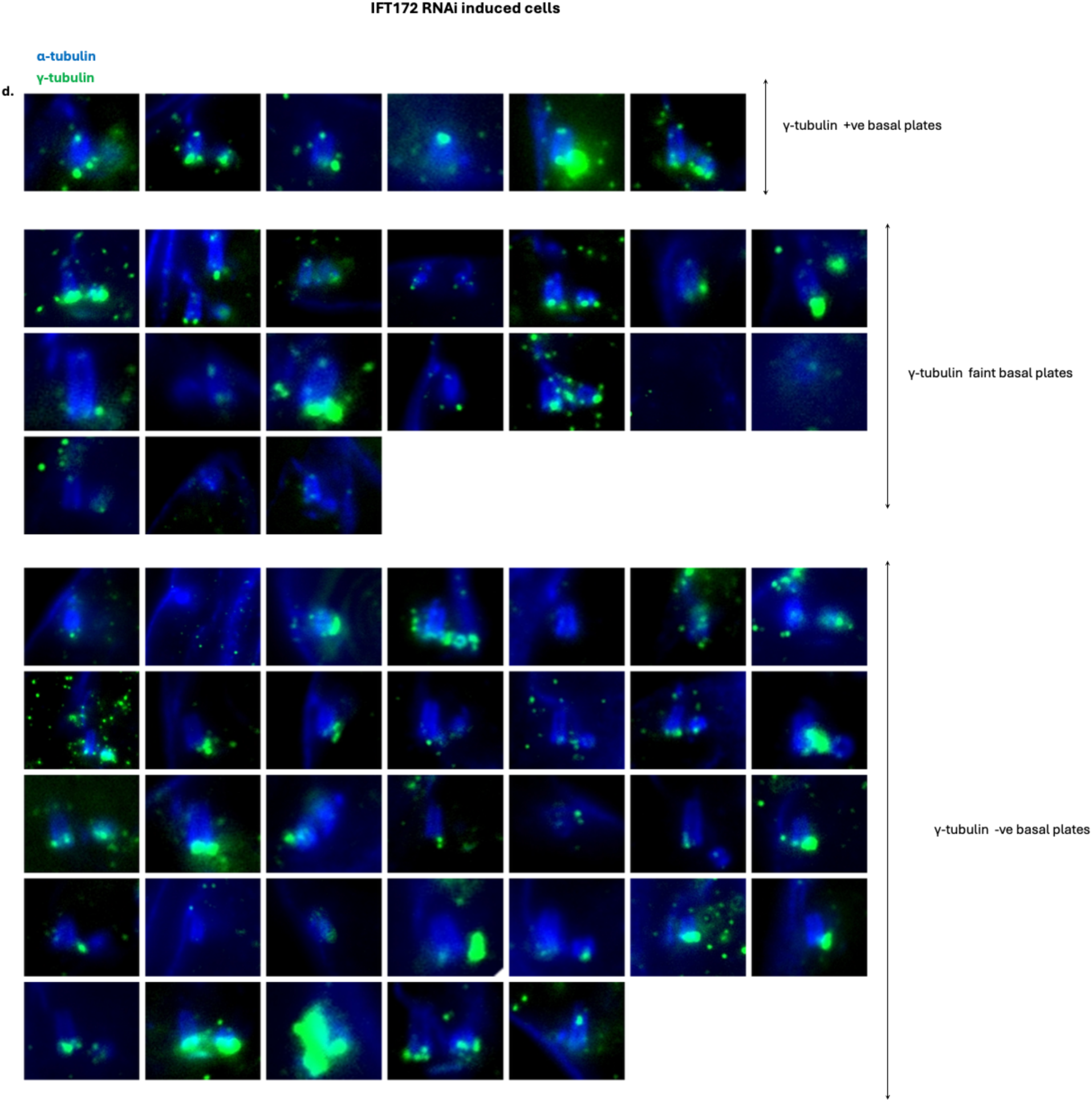
U-ExM images categorised for the presence/absence of γ- tubulin and TZP103.8 on the basal plate in control and IFT172 RNAi induced cells. a) TZP103.8 signal on the basal plate in control cells. b) γ-tubulin signal on the basal plate in control cels. c) TZP103.8 signal on the basal plate in IFT172 RNAi induced cells. d) γ-tubulin signal on the basal plate in IFT172 RNAi induced cells.

**Supplemental File 1.**
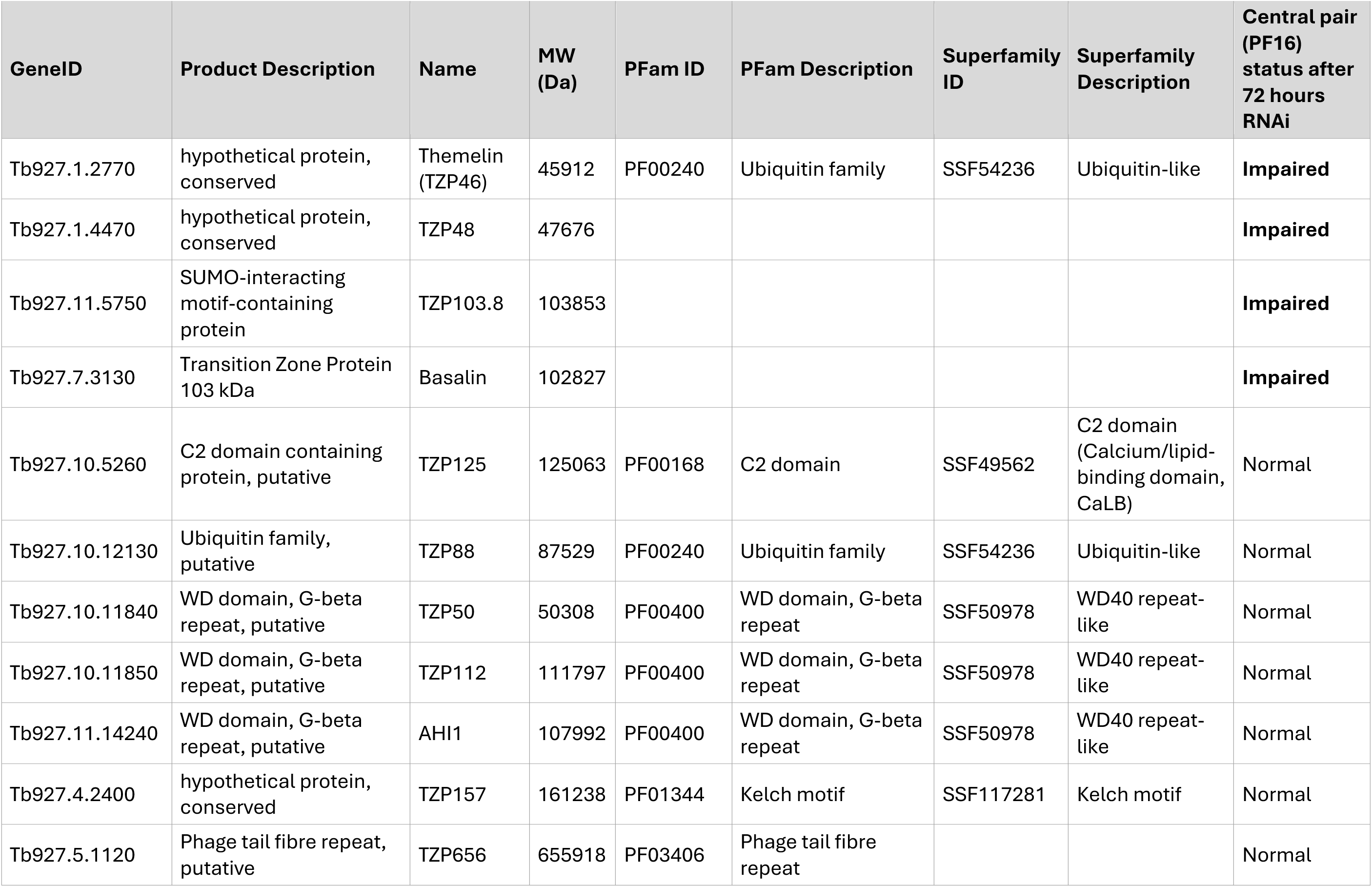
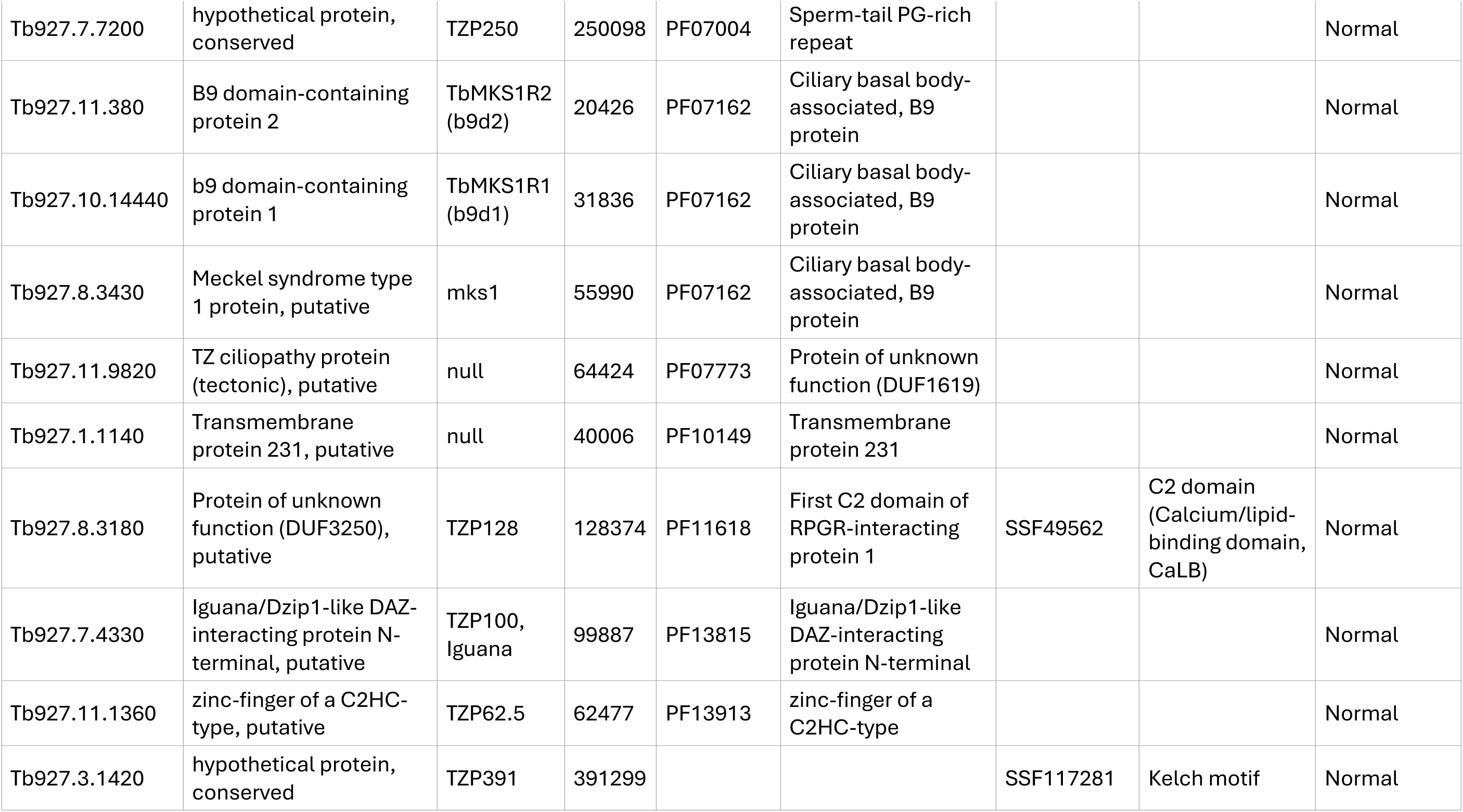
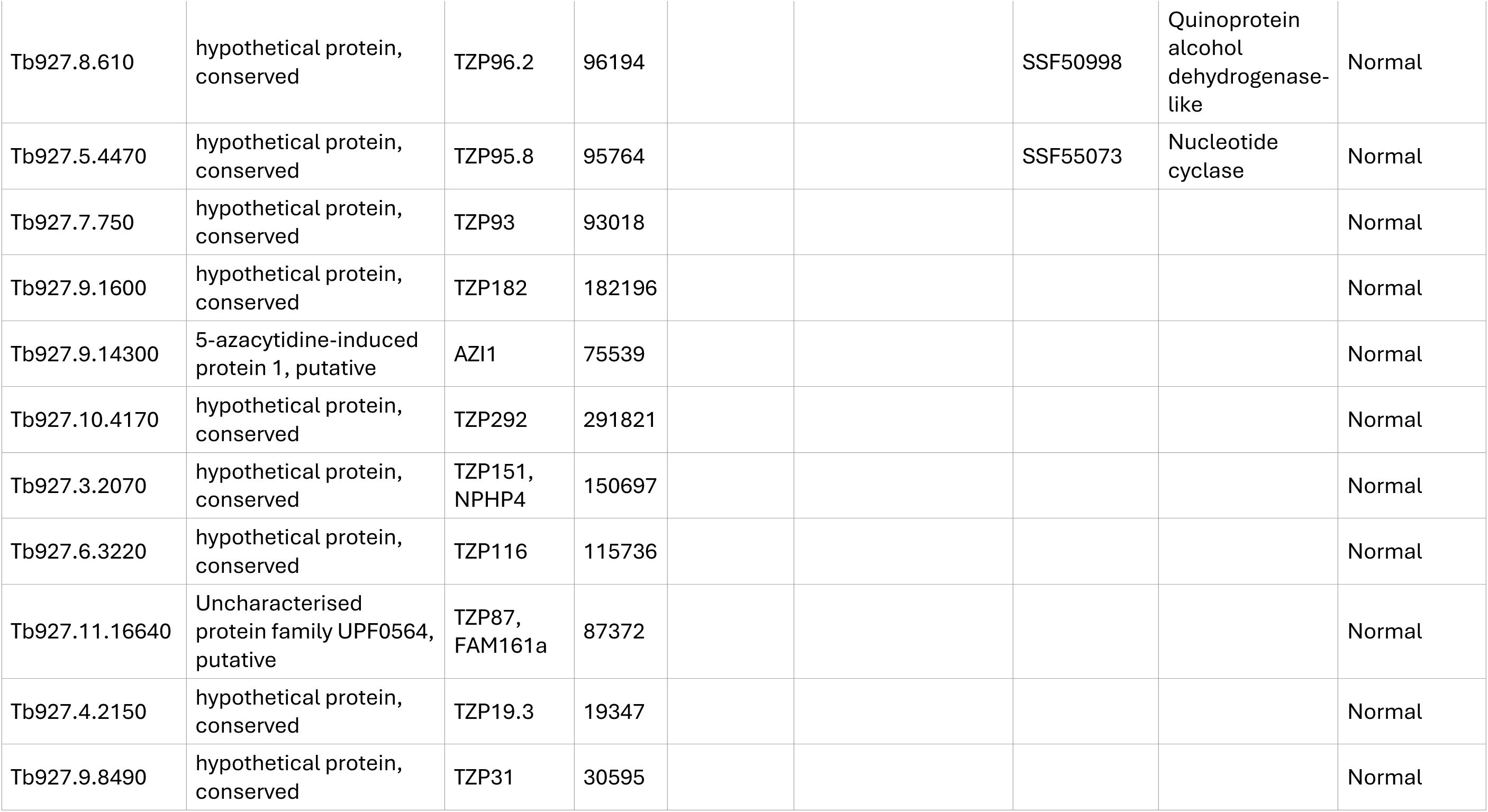
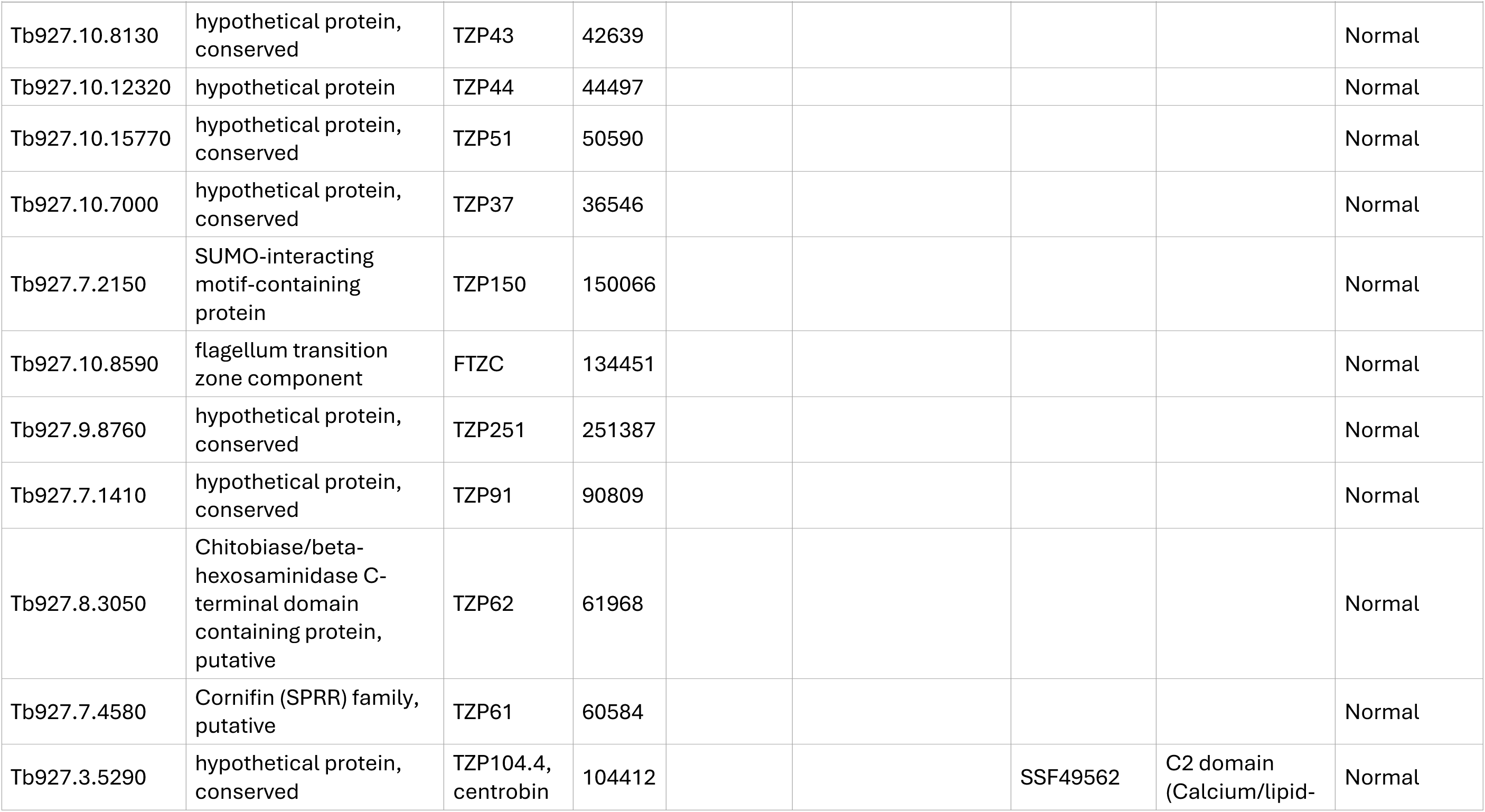
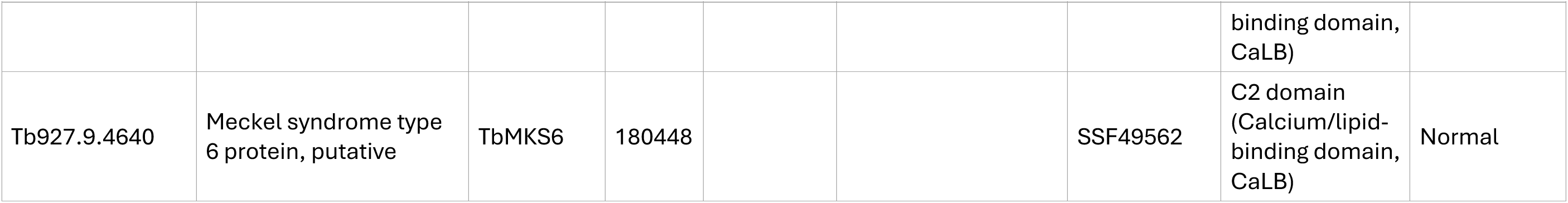
A list of TZPs target for RNAi and their biochemical attributes.

**Supplemental File 2.**
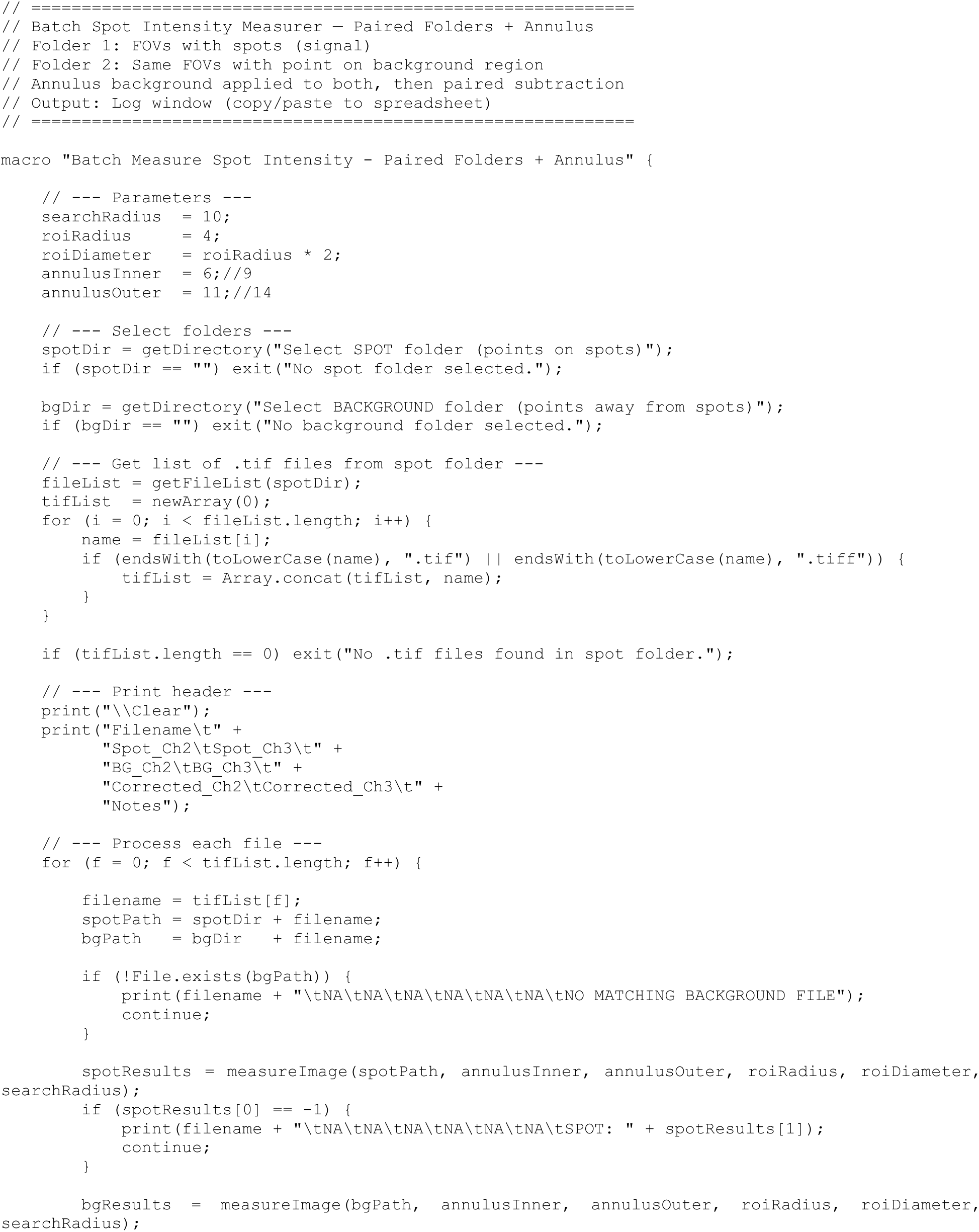
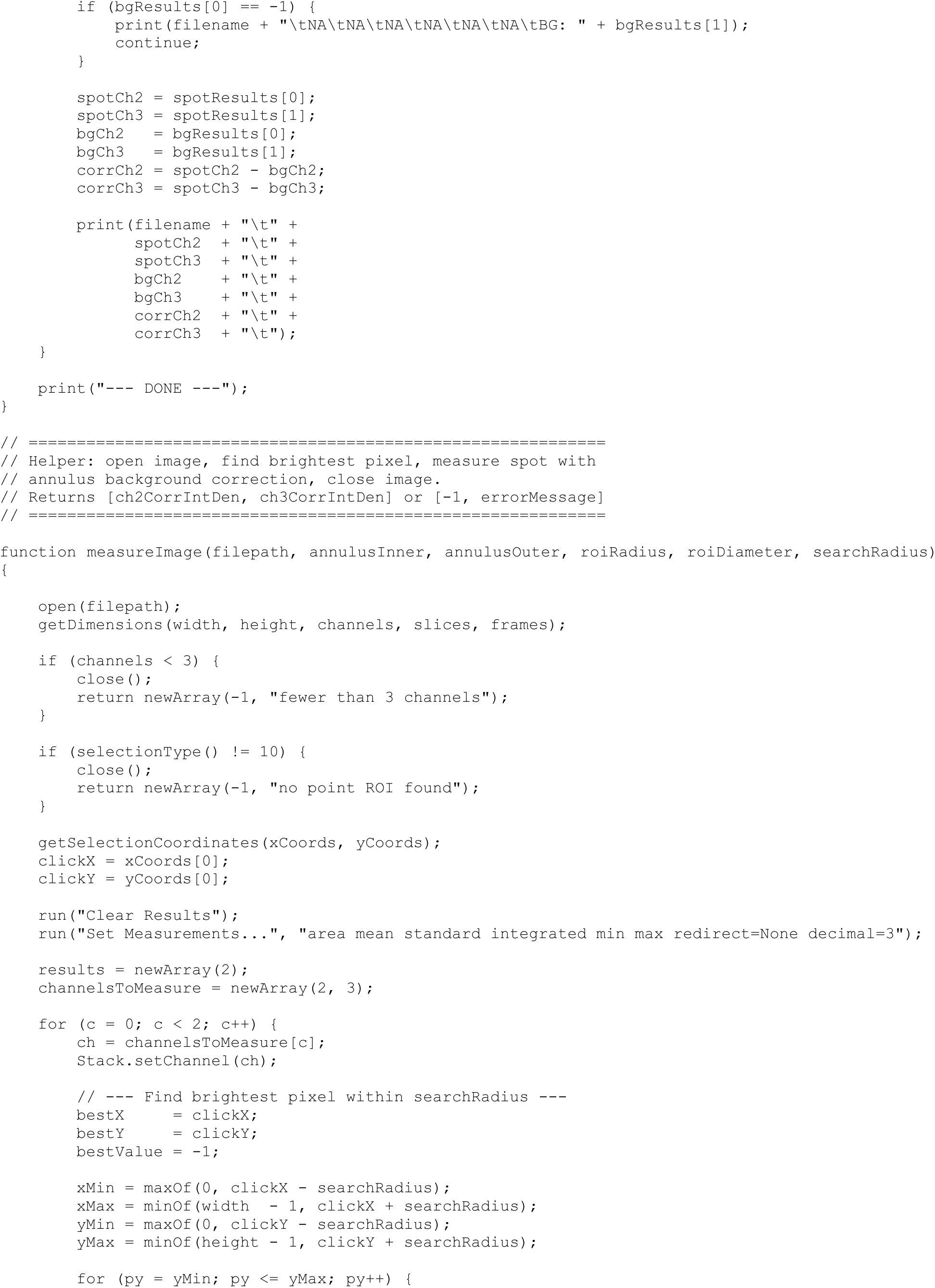
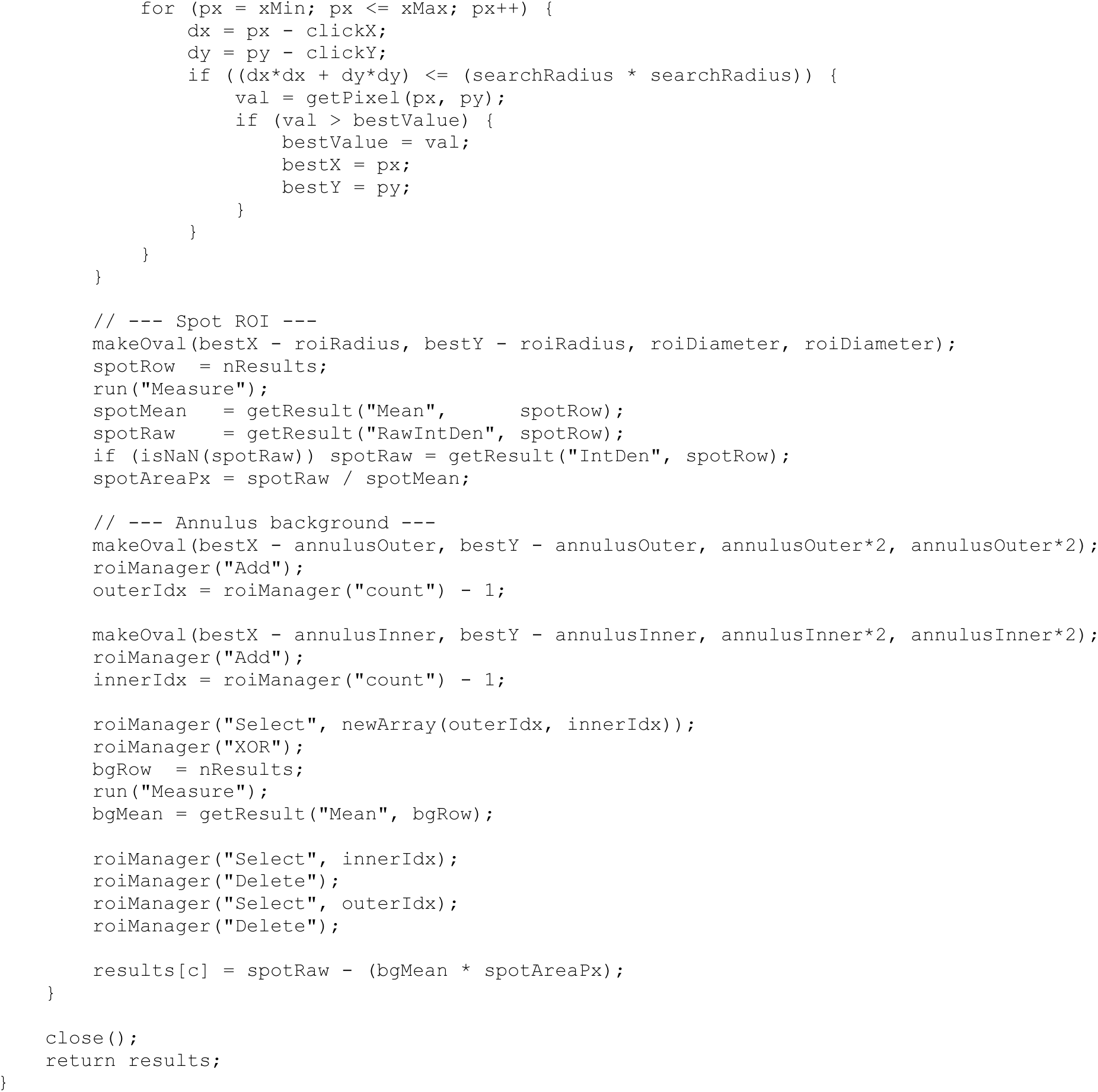
A FIJI macro for measuring TZ spot intensity.

